# The Effects of Lamin B Receptor knockdown on a Myeloid Leukemia Cell

**DOI:** 10.1101/2024.06.19.598074

**Authors:** David B. Mark Welch, Ada L. Olins, Donald E. Olins

## Abstract

In an effort to extend our understanding of the genetic functions of the nuclear envelope protein Lamin B Receptor (LBR), we examined the effect of a stable short hairpin (sh1) RNAi knockdown of LBR on the transcriptome and immunostained morphology of the human myeloid leukemia cell line (HL-60/S4). This examination was on sh1 cells induced to granulocytic form with Retinoic Acid (RA) versus sh1 cells maintained undifferentiated (0). By comparison to control cells (i.e., not sh1), we obtained gene lists that were differentially expressed only in the LBR knockdown cell line (i.e., “only-sh1-down” and “only-sh1-up”), in RA versus 0 cells. These curated gene lists were examined by Gene Ontology (GO) analysis. Aside from chromatin related GO terms, the most surprising finding was a significant downregulation of Golgi related genes only in the sh1 cells. Possible relationships between the “Cis-Golgi-Network” and LBR are discussed. Another surprise was a significant upregulation of “Ribosome” protein transcripts only in the sh1 cells. In parallel to these findings, an immunostaining comparison of nucleoli in S4 and sh1 cells demonstrated that the number and location of nucleoli in a single nucleus differs, depending upon the presence of LBR. Speculations on the influence of LBR levels upon the liquid-liquid phase separation model of nucleolar condensation are presented.

## Introduction

Lamin B Receptor (LBR) is a protein resident of the inner nuclear membrane (INM) in the interphase nuclear envelope (Giannios et al., 2017; Nikolakaki et al., 2017; Olins et al., 2010b). It was first described in the nuclear envelope of turkey erythrocytes (Worman et al., 1988). With subsequent characterization of human LBR (Ye and Worman, 1994), evidence was presented that the C-terminal 407 amino acids (aa) is embedded within the lipid-rich INM, organized into 7-8 transmembrane segments and possesses sterol reductase activity essential for cholesterol biosynthesis (Olins et al., 2010b; Tsai et al., 2016). The N-terminal 208 aa interacts with the lamina and underlying heterochromatin. Residues 1-60 aa constitute a Tudor Domain, which is believed to bind to the heterochromatic histone modification H4K20me2 (Hirano et al., 2012; Olins et al., 2010b). LBR residues 113-117 aa possess a Heterochromatin Protein 1 (HP1) Chromo Shadow Domain (CSD) binding motif “VxVxL”. The CSD connects to an HP1 Chromo Domain (CD), which binds to methylated lysine at H3K9 (Sanulli et al., 2019). It is generally accepted that LBR connects heterochromatin to the interphase nuclear envelope and also functions in sterol biosynthesis (Nikolakaki et al., 2017; Olins et al., 2010b). A previously published schematic version of LBR embedded within the INM, lamina and attached to heterochromatin is presented in Figure 1.

**Figure 1.**
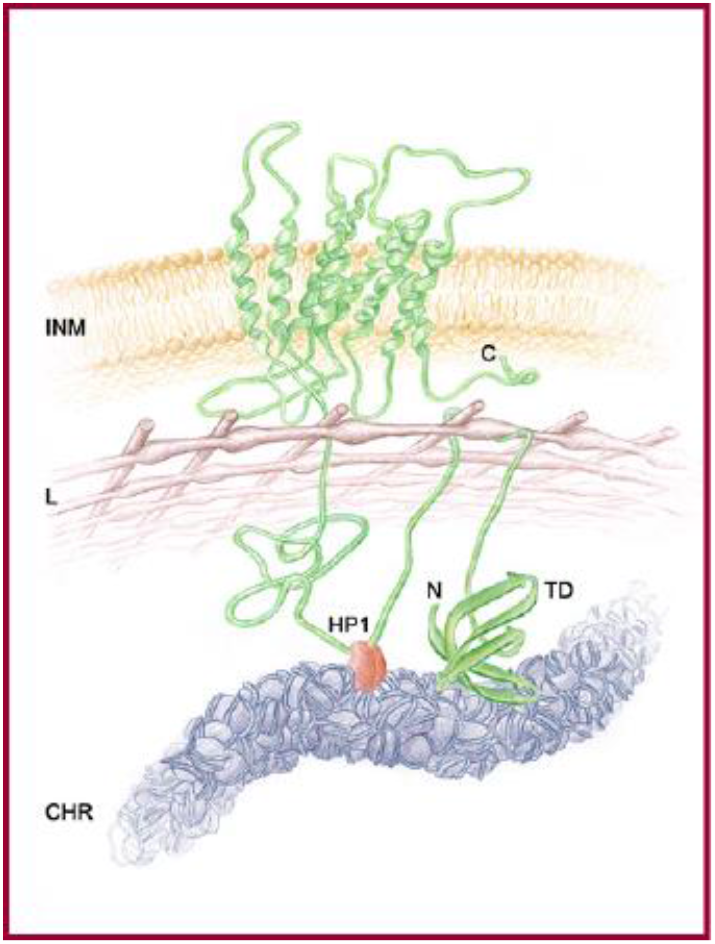
Schematic representation of LBR embedded within the nuclear envelope. Landmarks: INM, inner nuclear membrane (yellow); N and C, termini of LBR (green); L, lamina meshwork (brown); TD, Tudor domain; HP1, heterochromatin protein 1(red); CHR, peripheral heterochromatin with nucleosomes (blue). For illustrative purposes, the dimensions are distorted, with LBR drawn at a high magnification. Other structures are also not drawn to scale in this cartoon. For currently accepted dimensions: the INM is ∼5 nm thick; L, ∼15 nm thick; CHR, ∼20–30 nm diameter. This scheme was taken from (Olins et al., 2010b)

A variety of LBR mutations have been associated with human and mouse genetic diseases (Olins et al., 2010a; Olins et al., 2010b; Turner and Schlieker, 2016). At the benign level, heterozygous mutations in LBR result in the absence of nuclear lobulation of blood neutrophils (Pelger-Huet Anomaly). Homozygosity results in skeletal malformations and fetal death (Greenberg Skeletal Dysplasia [Chondrodysplasia] in humans; Ichthyosis in mice) (Greenberg et al., 1988; Hoffmann et al., 2002; Olins et al., 2010b; Shultz et al., 2003). This diversity of phenotypes resulting from LBR mutations has stimulated research analyses into the “multiple” genetic functions of LBR. Our approach has been to employ a Lentiviral “knockdown” of LBR in the human myeloid leukemia cell line (HL-60/S4). In this cell line, the immortalized promyelocyte can be differentiated *in vitro* into neutrophils with Retinoic acid (RA), into monocytes with Vitamin D and into macrophage with Phorbol Ester (TPA). The present study compares the polyA mRNA transcriptomes of the LBR mRNA knockdown cells (sh1) with two control cell lines, i.e., uninfected HL-60/S4 cells (S4) and lentiviral GFP-expressing HL-60/S4 cells (gfp). These three separate cell lines (sh1, S4 and gfp) are compared in the untreated (0 or “un”) state and in the treated (RA) state, with the purpose of discovering additional genetic functions of human LBR.

## Materials and Methods

### Cell Culture

HL-60/S4 cells (ATCC CRL-3306) were cultivated in standard RPMI-1640 medium plus 10% FCS and 1% Pen/Strep. Lentivirus derived HL-60/sh1 and /gfp cells were maintained in standard medium supplemented with 1mg/ml of puromycin (Olins et al., 2010a). Differentiation of sh1 and gfp cells with RA for 4 days, in the absence of puromycin, has been described earlier (Olins et al., 2010a).

### RNA purification

Quadruplicate samples (5×10^6^ cells/sample) of undifferentiated and retinoic acid (RA) treated HL-60/S4 cells, sh1 and gfp cells were centifuged, rapidly frozen and stored in liquid nitrogen (LN_2_). Samples were thawed by the addition of the RLT lysis buffer from the Qiagen RNeasy Mini Kit and RNA purified according to the manufacturer’s protocol. Care was taken to maintain RNAase-free conditions with RNAase Zap. RNA was eluted with molecular biology grade water, frozen in LN_2_ and shipped on dry ice to the Marshall University Genomics Core Facility. QC determinations (all 12 samples had a RIN score of 10), preparation of the libraries (Illumina mRNA library preparation Kit) and sequencing (Illumina HiSeq1500) was carried out at the Core Facility. The resulting sequences are available from the NCBI Sequence Read Archive as BioProject PRJNA674660.

### Data analyses and presentation

We analyzed the four libraries from each of the samples described above with previously generated data of undifferentiated and retinoic acid (RA) treated HL-60/S4 cells (Mark Welch et al 2017) available from NCBI Sequence Read Archive as BioProject PRJNA303179. We identified genes with significantly differential transcript levels following the RSEM-EBSeq workflow previously described (Mark Welch et al 2017, Mark Welch et al 2023) and outlined at http://deweylab.github.io/RSEM using the sequences and annotation of UCSC hg38. We used bowtie2 v2.3.2 to map paired-end reads to transcripts extracted from the reference genome, and calculated transcript level values using RSEM v1.3.0. RSEM uses a maximum likelihood expectation-maximization algorithm to estimate the transcript levels of isoforms from RNA-Seq reads (Li and Dewey, 2011). We then calculated the significance of relative expression differences using EBSeq v1.2.0 for gene-level analysis. EBSeq returns the normalized mean count of reads mapped using the median of ratios approach of DESeq (Anders and Huber, 2010) and the posterior probability of differential expression (PPDE) between conditions, which is naturally corrected for multiple tests (i.e., the PPDE is equivalent to one minus the false discovery rate, FDR). We conducted two 3-way tests: no knockdown (S4) vs LBR knockdown (sh1) vs gfp knockdown for untreated cells (0) and for RA treated cells (S4-RA vs sh1-RA vs gfp-RA). We conducted pairwise tests of each cell type with and without RA treatment (S4 vs S4-RA; sh1 vs sh1-RA; gfp vs gfp-RA). We accepted genes with PPDE>0.95 as having significantly different transcript levels. See Tables S1 and S2 for a complete list of EBseq normalized mean counts and PPDE values for each hg38 gene for 3-way and pairwise tests, respectively. Output from RSEM and EBSeq were imported into a MySQL database with hg38 annotation for analysis. We uploaded lists of differentially expressed genes (i.e., HGNC gene symbols) to WebGestalt (http://bioinfo.vanderbilt.edu/webgestalt) for over-representation analysis of GO non-redundant terms (Zhang et al., 2005). We considered a GO term over-represented if the hypergeometric test returned an adjusted p value (FDR) less than 0.05.

### Microscopy

Confocal imaging was performed on a Leica SP8 microscope. All images were collected as stacks, where each slice was 1024 x 1024 pixels. Z steps were 0.5µm. Confocal stacks were deconvolved with Autoquant X3 using an adaptive psf (point spread function). Figures were prepared using Fiji and Adobe Photoshop. Maximum intensity projections were prepared with Fiji (image/stacks//Z project). Rabbit (R), Mouse (M) and Goat (G) primary antibodies (employed at manufacturers’ recommended dilutions) were purchased from: Abcam R anti-LBR (ab32535), R anti-NPM1 (ab183340), R anti-TRIP11 (ab72223), R anti-H4 Multi-Ac (ab177790), M anti-GIANTIN (ab37266), G anti-HP1alpha/CBX5 (ab77256); Active Motif R anti-H3K9me2 (#39239) and R anti-H3K9me3 (#39161). Mouse anti-nucleosome acidic patch antibody (PL2-6) has been described earlier (Olins and Olins, 2018; Zhou et al., 2019). RNA specific staining employed SYTO RNASelect (ThermoFisher Scientific).

## Results

### The efficiency of LBR knockdown

Our previous electron microscopy results (Olins et al., 2010a), shown here (Figure 2), clearly demonstrated that LBR knockdown (employing a complementary short hairpin RNA, “sh1”) resulted in the <u>absence</u> of RA-induced neutrophil nuclear lobulation and nuclear envelope-limited chromatin sheets (ELCS) connecting nuclear lobes within one cell nucleus (Olins et al., 1998; Olins and Olins, 2009). We also demonstrated that knockdown of LBR mRNA in undifferentiated sh1 cells, resulted in decreased LBR protein, compared to S4 and gfp cells, as assayed by immunoblotting (Olins et al., 2010a). In the present study, we performed immunoblotting of LBR on S4 and sh1 cells, each following +/- RA treatment (Figure 3). The same number of cells (and amount of cell protein) were extracted in Laemmli sample buffer, as seen in the Coomassie stained left-half of the SDS PAGE. The PVDF membrane right-half of the PAGE gel is reacted with anti-LBR, with images collected at two different ECL exposure times. The relative reaction intensities are: S4 RA>S4 0>>sh1 RA>sh1 0. It is also clear that in the sh1 cells, with much reduced LBR, RA treatment produces a detectable increase in LBR.

**Figure 2.**
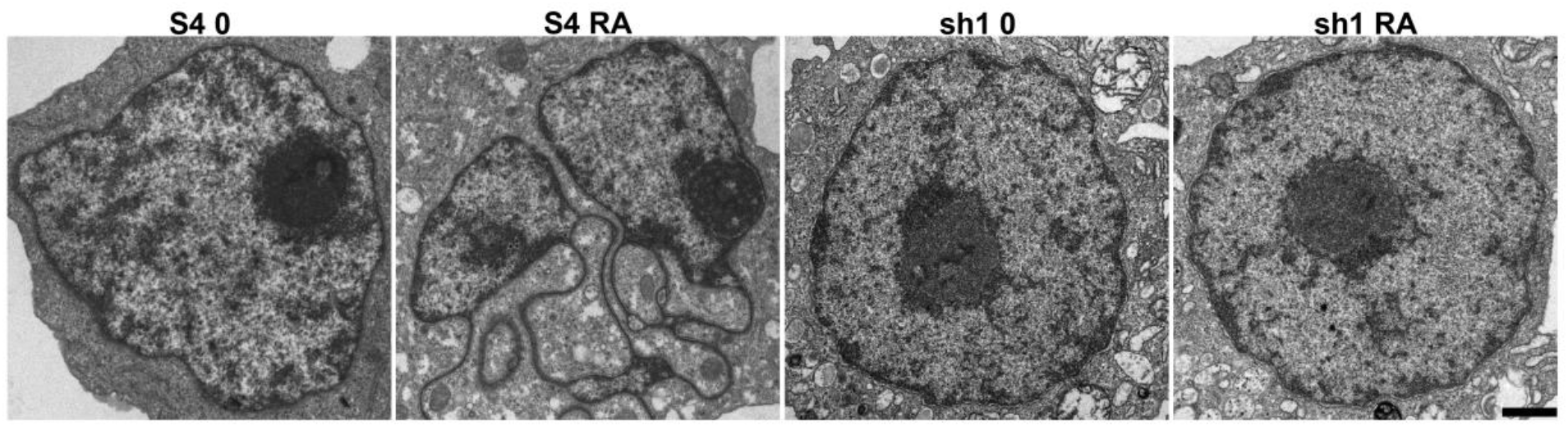
Thin-section electron microscopy of undifferentiated (0) and RA treated S4 and sh1 cells. Note the appearance of a lobulated nucleus and chromatin sheets (ELCS) connecting the nuclear lobes in the RA treated S4 cells (S4 RA) and their absence in the RA treated sh1 cells (sh1 RA). Note the positions of the densely stained nucleoli in the S4 and sh1 cells. Magnification bar: 1 μm. These images have been previously published (Olins et al., 2010a).

**Figure 3.**
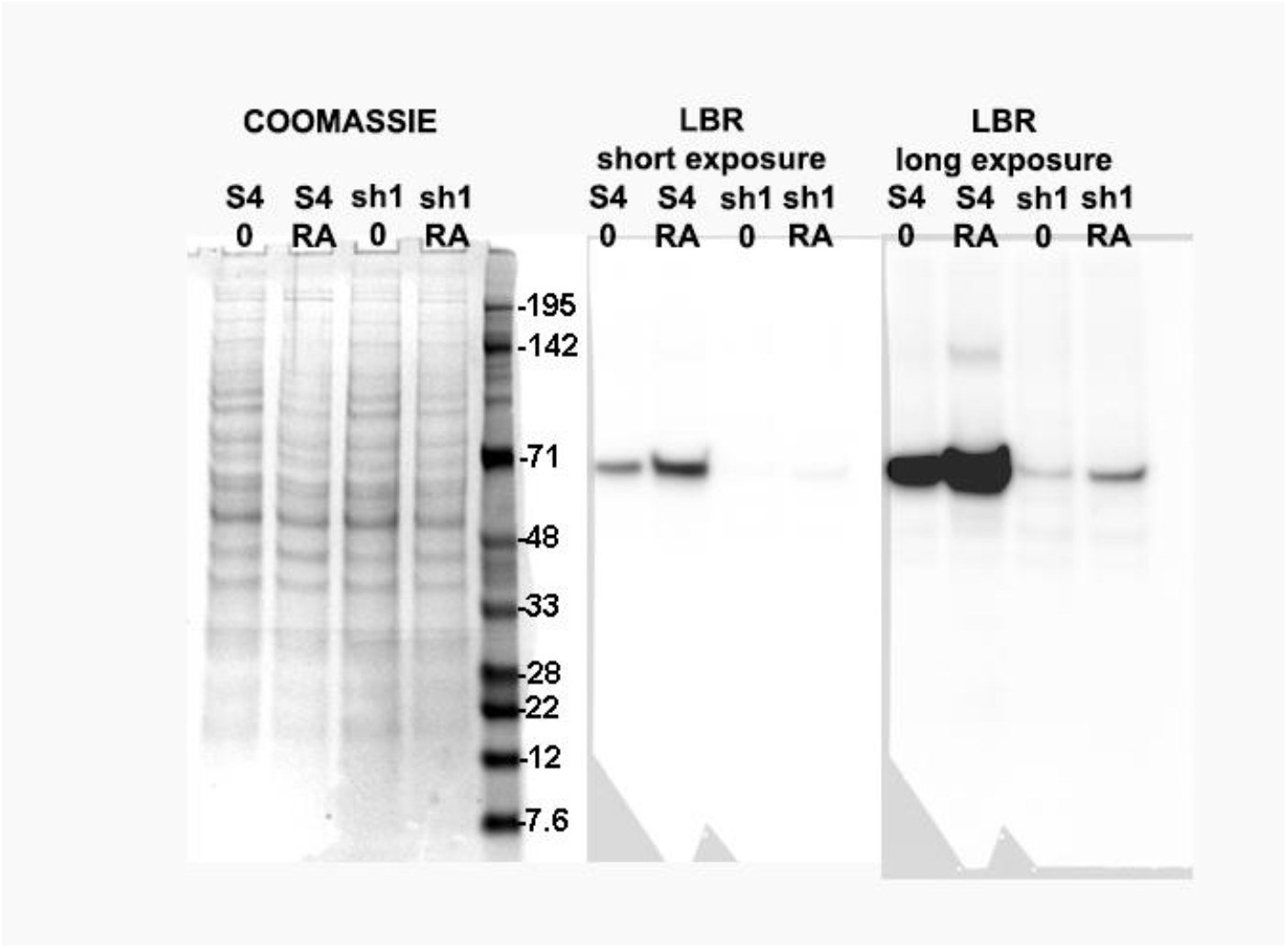
Immunoblot of LBR in S4 and sh1 cell extracts following (+/-) treatment with RA. Left half: Coomassie Blue stained gel adjacent to molecular weight (MW) markers (kDa). Right half: PVDF membrane with ECL reaction at short and long exposure times. Based upon the MW markers, the LBR band is ∼67 kDa.

S4 cells were cultured in standard RPMI-1640 medium without puromycin; sh1 and gfp cells were maintained in “selective” medium containing 1mg/ml of puromycin (Olins et al., 2010a). In the present study of the transcriptomes of sh1, S4 and gfp cells, sh1 cells exhibited a clear reduction (∼8 to 10-fold) in LBR polyA mRNA transcript levels, compared to S4 and to gfp cells (+/-RA) (Figure 4). For many of the subsequent analyses, comparisons are between sh1 and S4 cells. S4 and gfp cells are <u>not</u> identical “controls”. In this sense, we regard sh1, S4 and gfp cells as “separate” cell lines.

**Figure 4.**
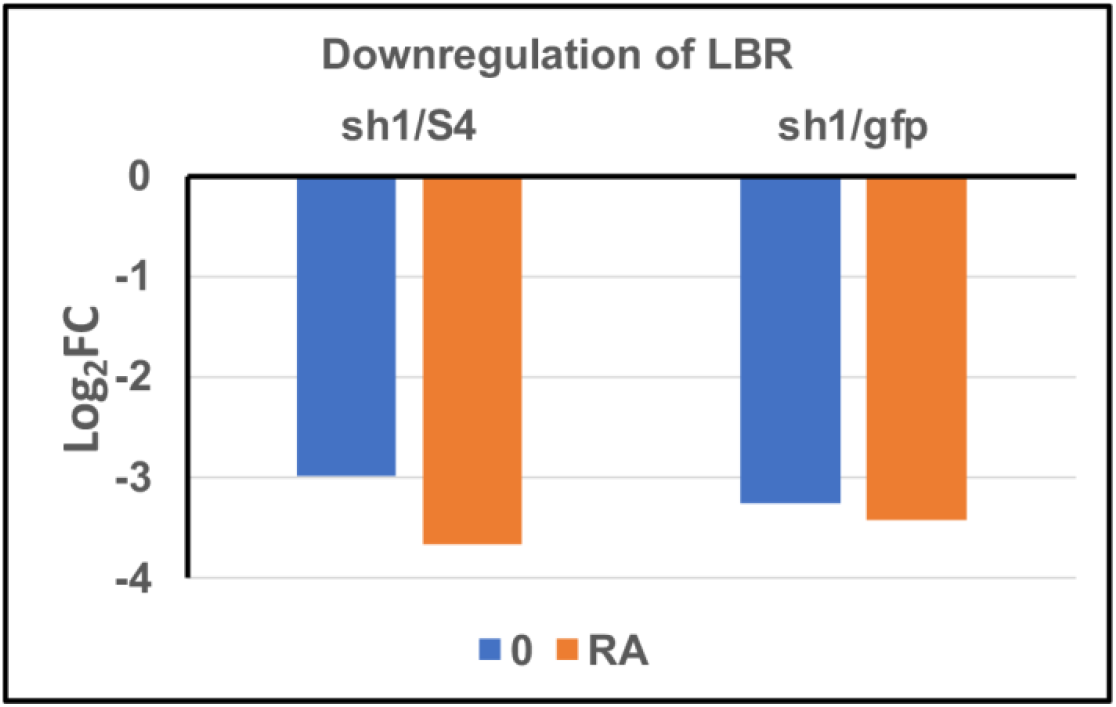
Downregulation (Log_2_ Fold Change<0) of LBR transcript levels, comparing sh1 with S4 cells (sh1/S4) and sh1 with gfp (sh1/gfp) cells, both for undifferentiated (0, blue) and for granulocytic (RA, orange) cell states. (Log2FC of -3 equals an 8-fold reduction). Graphs are derived from data in Table S1 “3-way”.

### Global transcriptome changes due to LBR knockdown

The goal of this study is to broadly identify LBR transcriptome knockdown effects on specific “functional” groups of gene transcripts. We searched for effects observed <u>only</u> in sh1 cells and <u>not present</u> in either S4 or gfp cells (+/-RA). The “only-sh1-down” gene list selects genes that are downregulated only in sh1 cells, not in S4 or gfp cells; the “only-sh1-up” list, features upregulated genes, only in sh1 cells, not in S4 or gfp cells. We employed these “curated” gene lists to identify specific Gene Ontology (GO) terms, where specific transcript levels are significantly increased (upregulated) or decreased (downregulated) in sh1 cells. Discovery of these GO term changes is intended to expand our understanding of the multiple functions of LBR, beyond those already observed in the microscopy experiments (Olins et al., 2010a).

The complete transcriptome data from sh1, S4 and gfp cells (+/-RA) are presented in Supplemental Tables S1 and S2. Table S1(a and b) “3-way” is useful for comparing the transcript levels among undifferentiated (denoted “00”) sh1, gfp and S4 cells (S1a) **or** among RA treated (“RA”) sh1, gfp and S4 cells (S1b). Table S2 “pairwise” presents **separate** cell lines (i.e., either sh1, gfp **or** S4) in pairwise comparisons. Each cell line is compared with and without RA, denoted “RA mean” and “00 mean” transcript levels, yielding Log (2) Fold Changes of RA/00. [The tabs and column headings are explained in the specific supplemental table legends].

Table 1 presents an overview comparison of Differential Gene Expression (DGE) for the three cell lines (sh1, S4 and gfp) responding to RA treatment for 4 days, employing the data derived from Table S2.

**Table 1.** Transcription level changes in HL-60/sh1, /gfp and /S4 cells after exposure to RA. DGE: Differential Gene Expression. Total: Total number of significant genes (PPDE≥ 0.95). # of genes: number of genes in each category of genes (Increased, Decreased, Unchanged). % of Total: percentage of genes in total number of significant genes. Only sh1: number of genes changed in sh1, but not in gfp and/or S4 cells. % of only sh1: percentage of “Only sh1” genes to “# of genes” (increased or decreased) in sh1 cells. Data derived from Table S2.

| RA/0 |  |  |  |  |  |  |  |  |  |
| --- | --- | --- | --- | --- | --- | --- | --- | --- | --- |
|  | sh1 |  |  |  |  | gfp |  | S4 |  |
| DGE | # of genes | % of total | Only sh1 | % of only sh1 | % of total | # of genes | % of total | # of genes | % of total |
| Increased | 4,134 | 24.6 | 696 | 16.8 | 4.1 | 4,469 | 26.6 | 3,989 | 23.7 |
| Decreased | 4,030 | 24 | 638 | 15.8 | 3.8 | 4,582 | 27.2 | 3,514 | 20.9 |
| Unchanged | 8,658 | 51.5 |  |  |  | 7,771 | 46.2 | 9,319 | 55.4 |
| Total | 16,822 |  |  |  |  | 16,822 |  | 16,822 |  |

Listed are the number of categorized genes exhibiting significantly increased transcript levels (“upregulation”), decreased transcript levels (“downregulation”) and unchanged transcript levels for each cell line. Further displayed are the % of genes (within each category), calculated by comparison to the total number of statistically significant genes (16,822) in the transcriptome dataset. Several observations are immediately apparent: 1) The number and % of increased (upregulated) genes comparing among the three cell lines are approximately similar to one-another. Likewise, for a comparison among decreased (downregulated) genes, and for a comparison among unchanged genes. It appears that RA treatment may have similar quantitative effects upon the majority of genes in the three cell lines. 2) Restricting the gene counting to “Only-sh1” cells (i.e., to upregulated genes (“only-sh1-up”) or downregulated genes (“only-sh1-down”) results in a considerable reduction of the number of genes available for GO analysis. Indeed, only 4.1% (696) of the total number of significant genes (16,822) are specifically upregulated in sh1 cells following RA treatment; only 3.8% (638) are specifically downregulated in sh1 cells following RA treatment. These two curated gene lists were employed in the GO analyses, focused upon the “Cellular Component noRedundant” function database; see below (*Gene Ontology Analysis)*.

### LBR and Nuclear Envelope Structure

The interphase nuclear envelope (NE) contains many proteins fulfilling numerous cellular functions (e.g., stabilizing the NE, attaching heterochromatin to the NE, connecting to the cytoskeleton, establishing pores in the NE etc.). Figure 5 demonstrates that the transcript levels for some of these NE proteins change during granulocyte differentiation induced by RA treatment, and that the <u>direction</u> of change (i.e., up- or downregulation) is generally the same for any specific gene, comparing sh1 and S4 cells.

**Figure 5.**
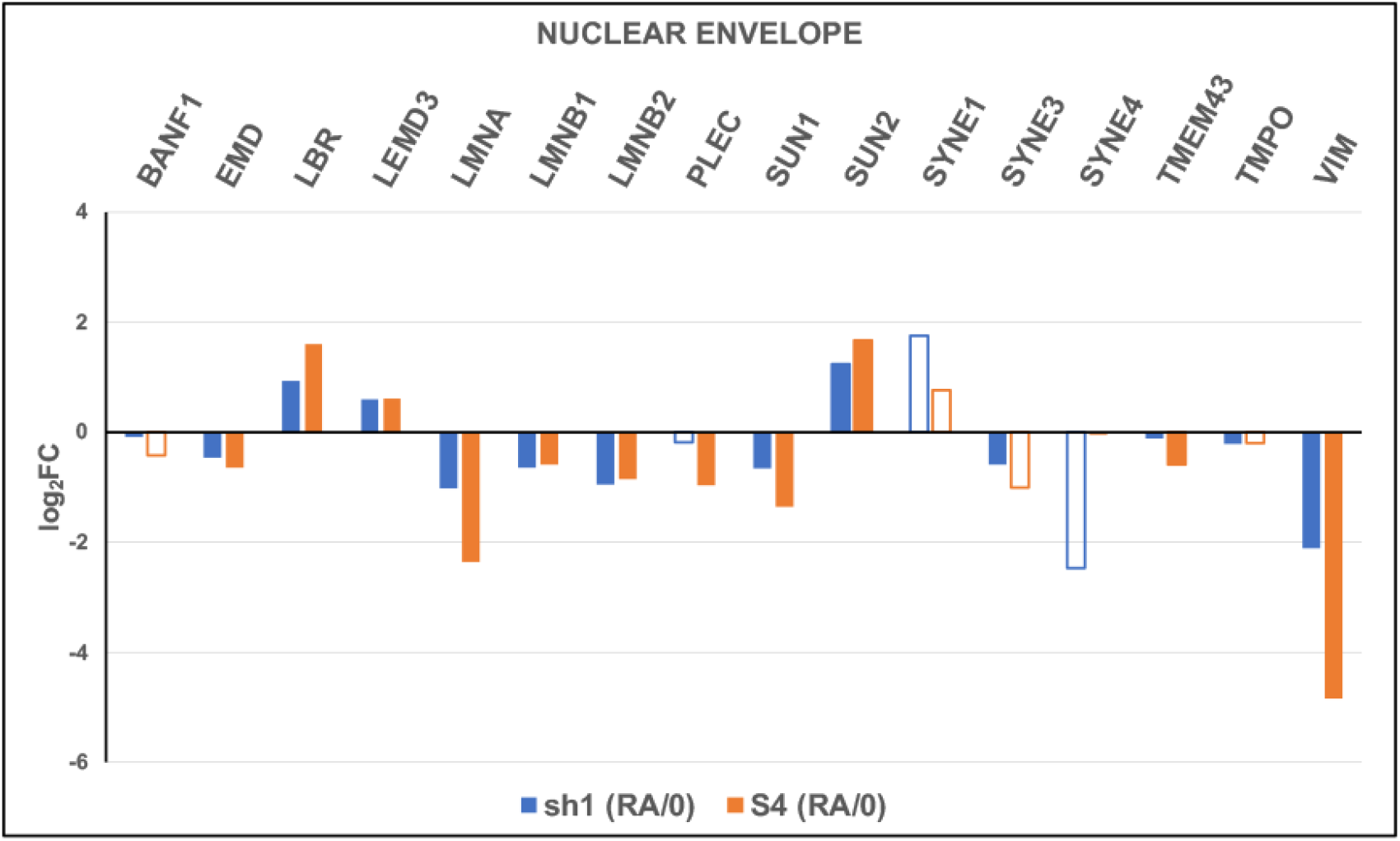
Log_2_FC changes of nuclear envelope (NE) protein transcript levels, comparing RA-treated with untreated (0) cells (RA/0). Generally, specific gene transcript levels responded in parallel directions (up-or-down) in both sh1 (blue) and S4 (orange) cells. Although the LBR transcript levels in sh1 and S4 cells are different, the Log_2_FC changes are in the same direction. Closed bars: PPDE>0.95 (significant data). Open bars: PPDE<0.95 (not significant). VIM (vimentin) resides in the cytoplasm but is connected to the NE protein PLEC (plectin). Plots are derived from data in Table S2 “pairwise”. HGNC Gene codes are displayed above the relative transcript level bars.

As described earlier, LBR functions as an essential sterol reductase in the biosynthetic pathway leading to cholesterol. Ironically, the LBR promoter does <u>not</u> possess a “sterol response element (SRE)”, unlike many of the other genes in the cholesterol biosynthesis pathway (Clayton et al., 2010; Cohen et al., 2008; Malu et al., 2016; Subramanian et al., 2012; Tsai et al., 2016; Turner and Schlieker, 2016). LBR resides in the interphase INM, an extension of the endoplasmic reticulum (ER), which possesses most of the other enzymes of the cholesterol biosynthesis pathway (Simons and Ikonen, 2000).

It is not clear by what mechanisms RA increases the level of LBR transcripts and whether RA (via a RA receptor protein) binds directly to the LBR promoter (Malu et al., 2016). After 4 days of RA-induced granulocyte differentiation in HL-60/S4 cells, the LBR transcript level is increased ∼3-fold (Mark Welch et al., 2017) and the nucleus is lobulated with the formation of extensive ELCS (Figure 2). Immunoblots of extracts from the same number of untreated and of (4 day) RA differentiated HL-60/S4 cells indicate that LBR protein is increased ∼3-4 fold (Olins et al., 2000); see Figure 3. It is also not understood how increased LBR levels result in increased granulocyte nuclear lobulations. In addition, a linear positive correlation of the number of LBR gene copies and the number of neutrophil nuclear lobes has been dramatically demonstrated (Gravemann et al., 2010). The underlying supposition of the role of LBR in the formation of nuclear lobes and ELCS is that increased levels of LBR result in localized INM growth (presumably, by virtue of increased local cholesterol concentrations) combined with <u>nuclear invaginations</u> of the “spreading” INM. It is proposed that as the INM grows, heterochromatin is attached to the increased membrane surface area, employing the elevated levels of LBR (Olins and Olins, 2009). Furthermore, ELCS have a remarkable uniformity of thickness, appearing to be two “face-to-face” INMs, each associated with one sheet of parallel ∼30 nm chromatin fibers forming a double-layer chromatin “sandwich” of ∼60 nm thickness (in the frozen hydrated form) (Xu et al., 2021).

### LBR, Cholesterol Biosynthesis & Retinoic Acid

Continuous RA (1mM) treatment of undifferentiated HL-60/S4 cells for 4 days results in upregulation (relative to untreated cells) of most gene transcripts involved with cholesterol biosynthesis (Mark Welch et al., 2017). Surprisingly, LBR knockdown in <u>untreated</u> sh1 cells also results in <u>upregulation</u> of cholesterol biosynthesis transcripts relative to the same transcripts in undifferentiated S4 cells (Figure 6; derived from Table S1 “3-way”). Note that the LBR transcript level is strongly reduced in sh1 cells, relative to S4 (as described earlier, Figure 4). Among the relatively upregulated genes is TM7SF2, which also functions as a C-14 sterol reductase, similar to LBR (Bennati et al., 2006; Lemmens et al., 1998). It should also be noted that sh1 cells exhibit “vigorous growth” and maintain the capacity for cell differentiation (Olins et al., 2010a). The only obvious difference is that sh1 cells do not exhibit nuclear lobulation and ELCS following RA treatment (Olins et al., 2010a). It is conceivable that TM7SF2 compensates for the knockdown of LBR, permitting sterol biosynthesis to proceed sufficiently for cell needs. Figure 6 also demonstrates that RA treated sh1 cells do not show the same degree of upregulation in most cholesterol biosynthetic genes, as observed in undifferentiated sh1 cells. Presumably, RA treated sh1 cells have their cholesterol needs satiated, with the help of TM7SF2.

**Figure 6.**
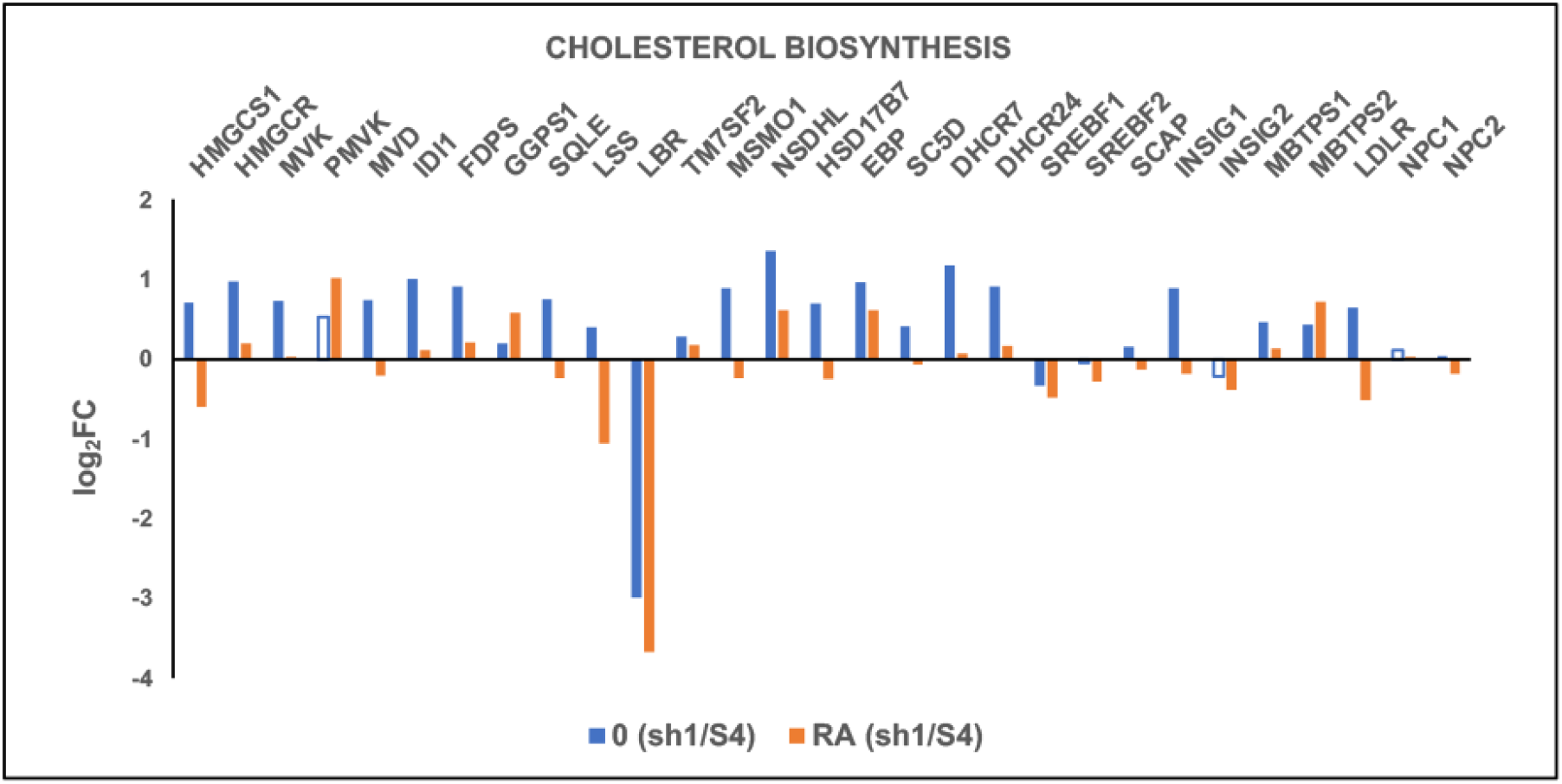
Relative transcript levels (Log_2_FC) for cholesterol biosynthesis genes, comparing sh1 to S4 cells for undifferentiated cells (0, blue) and for granulocyte cells (RA, orange). Note that the majority of genes in untreated 0 cells display upregulation, which is less consistent in treated RA cells. Closed bars: PPDE>0.95 (significant data). Open bars: PPDE<0.95 (not significant). Plots are derived from data in Table S1 “3-way”. HGNC Gene codes are displayed above the relative transcript level bars.

The observation that RA treated sh1 cells are not multi-lobulated opens the question of whether these cells are actually “granulocytes”, as are RA treated S4 cells. To examine this question, we compared the transcript levels of sh1 and S4 for identified “cytoplasmic granular proteins” (Mark Welch et al., 2017). Figure 7 compares the relative transcript levels (Log_2_FC) of treated sh1 and S4 cells (RA/0) for selected granular protein transcripts. It is immediately apparent that these transcription bars for each of the selected genes demonstrate parallel changes (up-or-downregulation), supporting that <u>both</u> cell types (sh1 and S4) are granulocytes. The magnitude of changes of S4 cells was uniformly “greater” than for sh1, suggesting that granulocyte differentiation of S4 cells may be “more complete” than for sh1 cells. The origin and significance of this possible difference in magnitude of differentiation is presently unknown, but likely depends on the level of LBR

**Figure 7.**
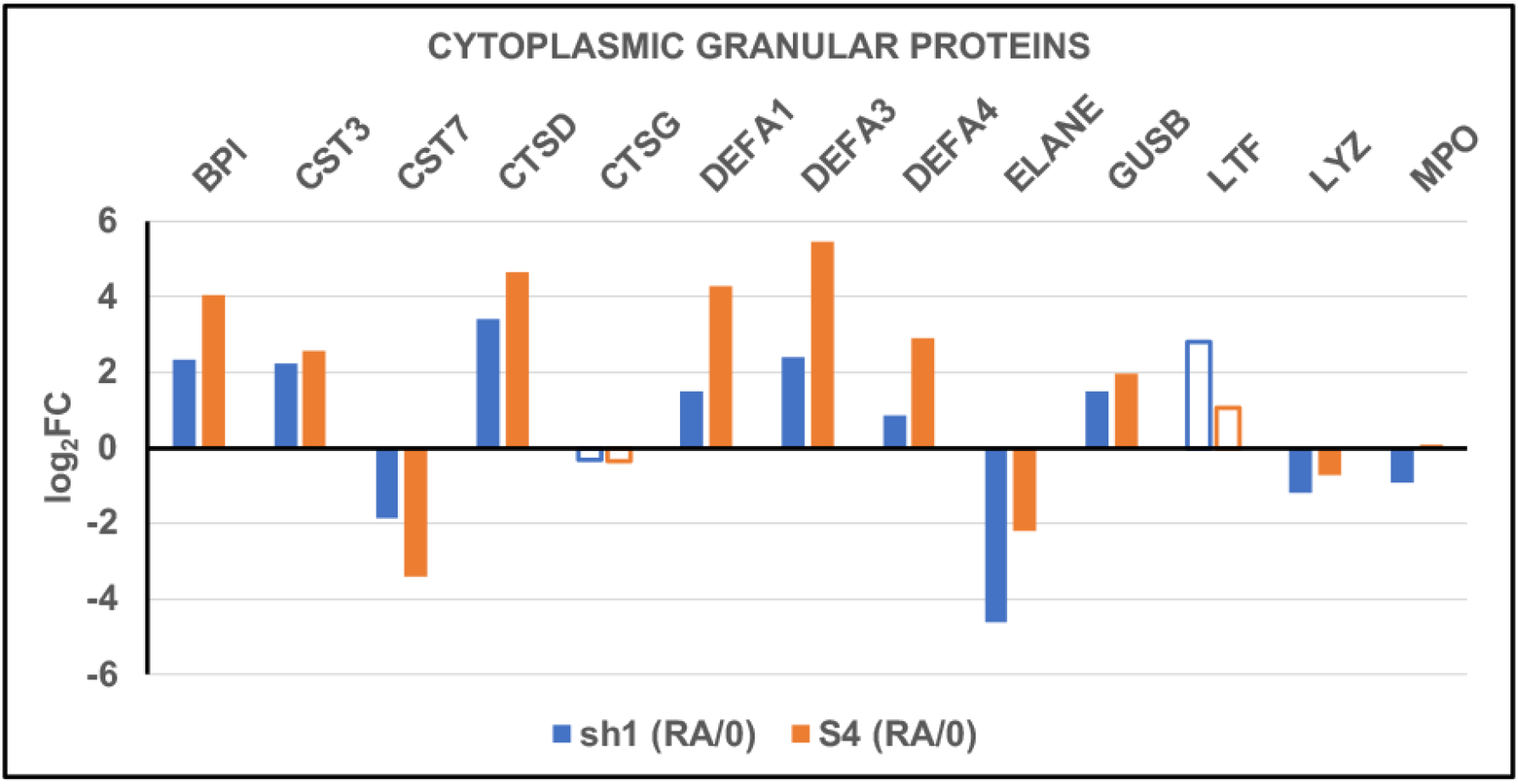
Relative transcript levels (Log_2_FC) for cytoplasmic granular proteins genes, comparing the transcript levels of RA treated sh1 and S4 cells to untreated cells (RA/0). Note that the individual gene transcripts display parallel up-or-down changes. Plots are derived from data in Table S2 “pairwise”. HGNC Gene codes are displayed above the relative transcript level bars.

### Gene Ontology Analysis-The Golgi Apparatus

Table 2 presents the “top 10” GO terms for the curated gene lists “only-sh1-down” and “only-sh1-up” derived from Table S2 “pairwise”. For us, GO was a true “discovery tool”. At the top of the “only-sh1-down” list, with FDR≤0.05 is (GO:0005801) “Cis-Golgi Network”. We had expected GO terms relevant to the nuclear envelope and chromatin, which are also seen (GO:0000793) “Condensed Chromosome” and (GO:0000151) “Chromosomal Region”, but <u>not</u> a GO term relevant to the Golgi apparatus.

**Table 2.** GO Terms.

| <u>Only-sh1-down</u> | <u>638 genes</u> | <u>Only-sh1-up</u> | <u>696 genes</u> |
| --- | --- | --- | --- |
| **FDR ≤0.05** |  | **FDR ≤0.05** |  |
| GO:0005801 | Cis-Golgi Network | GO:0005840 | Ribosome |
| GO:1902493 | Acetyltransferase Complex | **FDR>0.05** |  |
| GO:0034708 | Methyltransferase Complex | GO:0036452 | ESCRT Complex |
| GO:0030496 | Midbody | GO:0032994 | Protein-Lipid Complex |
| GO:0016607 | Nuclear Speck | GO:0017053 | Transcription Repressor Complex |
| GO:0000793 | Condensed Chromosome | GO:0005844 | Polysome |
| GO:0000151 | Ubiquitin Ligase Complex | GO:0031091 | Platelet Alpha Granule |
| GO:0098687 | Chromosomal Region | GO:0042579 | Microbody |
| **FDR>0.05** |  | GO:0044445 | Cytosolic Part |
| GO:1904949 | ATPase Complex | GO:0005875 | Microtubule Associated Complex |
| GO:0030139 | Endocytic Vesicle | GO:0005667 | Transcription Factor Complex |

The Golgi apparatus is aptly described as “a processing and sorting hub in the transport and targeting of soluble cargo proteins and lipids to different destinations in the cell” (Liu et al., 2021). The “Cis-Golgi Network” (CGN)** consists of cisternae (i.e., membrane sacs) facing the ER (endoplasmic reticulum). Newly synthesized proteins in the ER are packaged into vesicles which travel to the CGN for incorporation into proximal cisternae. These proteins migrate down through the stack of cisternae, undergoing modifications (e.g., glycan processing), finally emerging as vesicles from the “Trans-Golgi Network (TGN), some as secretory granules fusing with the plasma membrane followed by exocytosis. Many of the Golgi proteins (e.g., “Golgins”) act to “catch” (tether) the vesicles traveling in the cytoplasm (Lowe, 2019). There are also reports of direct contact between Golgi cisternae and intracellular membranes (David et al., 2021), possibly “by-passing” vesicle formation and fusion.

The gene dataset for the GO term (Cis-Golgi Network) lists 15 genes, most of which are clearly Log_2_FC downregulated in sh1 cells, compared to transcript levels in S4 cells (Figure 8). Because of future discussion, we call particular attention to Golgin TRIP11, which is considerably reduced in sh1 (RA/0) cells compared to S4 (RA/0) cells.

**Figure 8.**
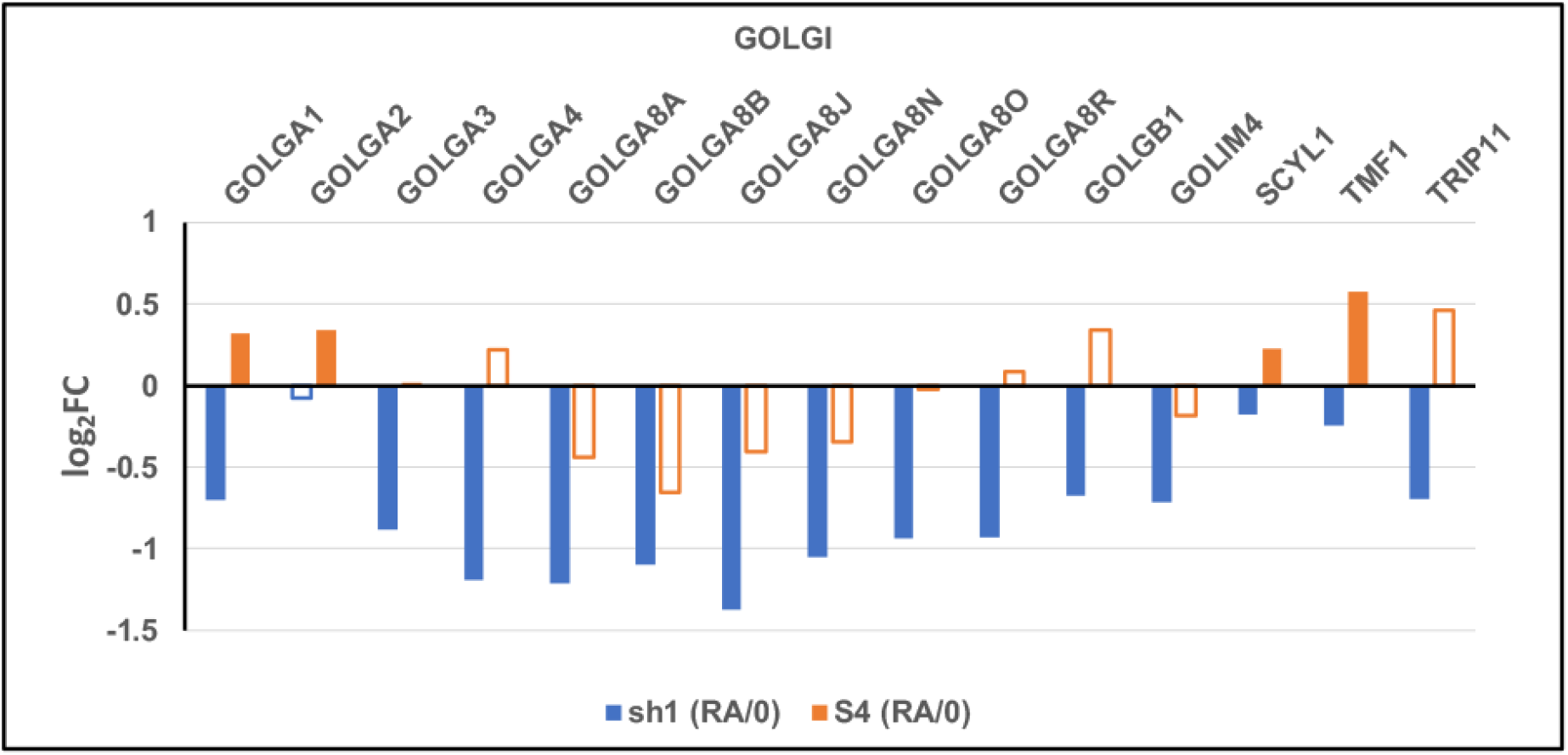
Relative transcript levels (Log_2_FC) for Cis-Golgi Network proteins: (**Blue** bars) Comparing sh1 cells (RA/0); (**Orange** bars) Comparing S4 cells (RA/0). Note that TRIP11 is downregulated in sh1 cells more than in S4 cells. Closed bars: PPDE>0.95 (significant data). Open bars: PPDE<0.95 (not significant). Plots are derived from data in Table S2 “pairwise”. HGNC Gene codes are displayed above the relative transcript level bars.

Regarding the Cis-Golgi Network, the principal question is: How does a significant reduction in LBR and/or sterol biosynthesis produce a “disabled” functioning Golgi apparatus? Most (60-90%) of cellular cholesterol is in the plasma membrane. The ER contains only ∼0.5-1.0%. The Golgi apparatus is intermediate in cholesterol content between these two types of membranes (Luo et al., 2017). Membrane sterol content increases progressively through the (Golgi-associated) secretory pathway (Bankaitis et al., 2012). We speculate that LBR knockdown with possible reduced levels of membrane cholesterol might decrease the efficiency of the Golgi apparatus in tethering vesicles, in sorting, in protein glycosylation, and/or cellular secretion.

We scoured the scientific literature for studies relating LBR to the Golgi and found one recent important publication (Wehrle et al., 2018). This study is concerned with the pathogenesis of a human chondrodysplasia (ACG1A) resulting from mutations in the Golgi protein (TRIP11). The authors were interested in a phenocopy of ACG1A, observed with Greenberg Dysplasia, a chondrodysplasia arising from LBR mutations (Greenberg et al., 1988). They performed RT-PCR on primary fibroblasts from a confirmed ACG1A fetus, which demonstrated reduced LBR mRNA levels compared to control fetal fibroblasts. Employing both immunofluorescent and electron microscope studies, they concluded that the TRIP11 mutation and the LBR mutation each affect Golgi architecture and functioning. They proposed that either of these mutated genes result in Golgi fragmentation, impaired vesicular trafficking, reduced cargo secretion and altered protein glycosylation. Our transcriptome data (Figure 8) clearly demonstrates that LBR knockdown in HL-60/sh1 cells leads to mRNA downregulation of a large set of Golgi genes, including TRIP11 (gene code: GMAP-210) and Giantin (gene code: GOLGB1), both “Golgi markers”.

We performed immunofluorescence (IF) microscopy (Figure 9) on S4 and sh1 cells, both undifferentiated (0) and differentiated (RA) cell forms, employing antibodies against TRIP11 and Giantin. The IF results, although subtle, generally agree with the transcriptome data (Figure 8). The IF images (Figure 9) are presented as Maximum Intensity Projections (MIP). Comparing granulocytic (RA-treated) cell forms, S4 cells revealed numerous “fine specks” of TRIP11 staining outside the boundaries of the Giantin-stained Golgi apparatus. These “fine specks” are almost completely absent in granulocytic sh1 cells. The specks could represent the described Golgi vesicles (Wehrle et al., 2018). Similar specks are seen around mitotic chromosomes, probably due to Golgi fragmentation during mitosis. Some staining appears to be within nuclei or chromosomes, due to MIP through over-and-under layers of cytoplasm.

**Figure 9.**
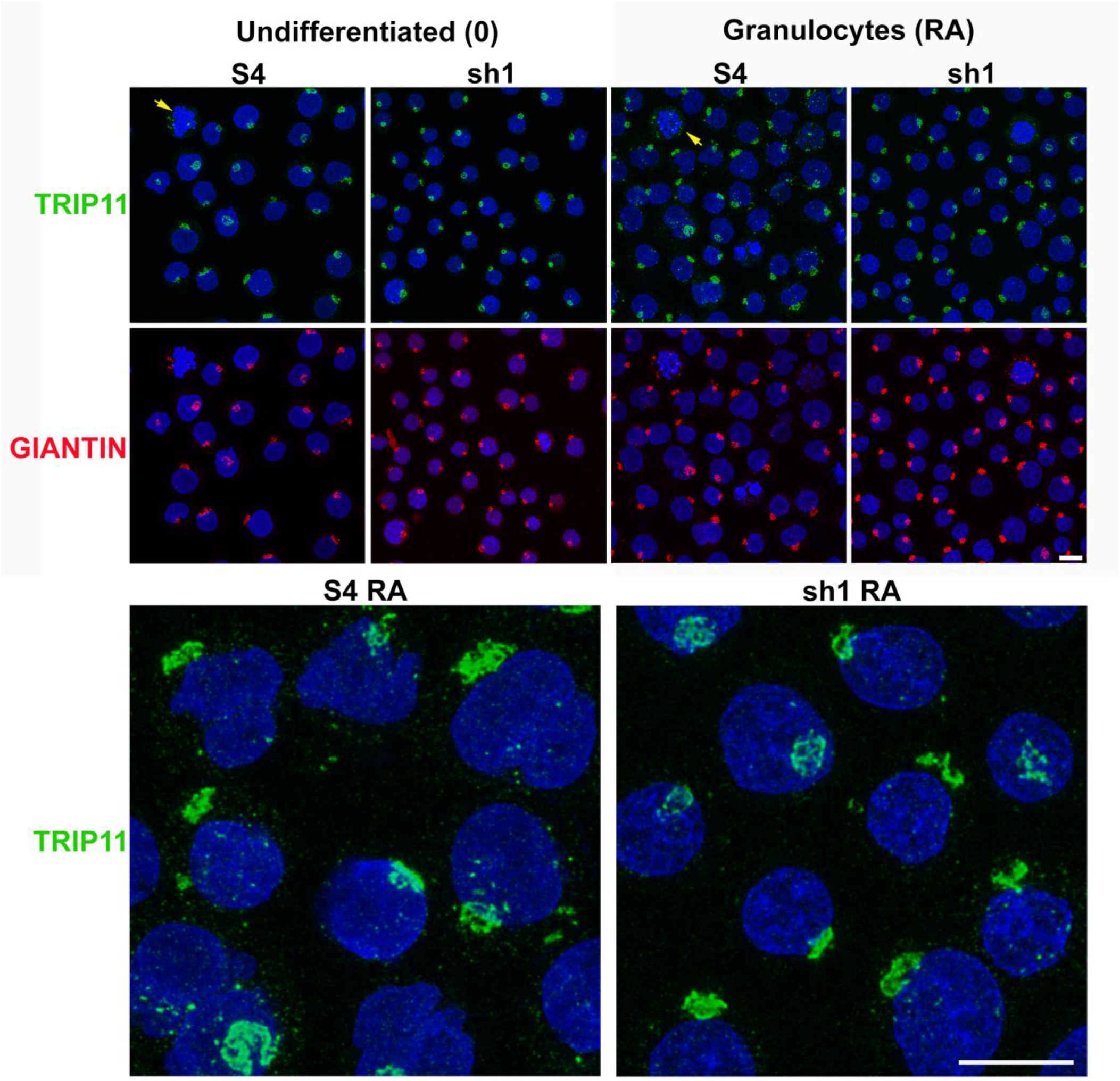
Immunostaining of the Golgi apparatus with anti-TRIP11 (green) and anti-Giantin (red). The top row and bottom row (five-fold enlargements of selected granulocyte regions in the top row) display TRIP11 staining. The middle row displays Giantin staining of identical fields presented in the top row. Nuclei and chromosomes are stained with DAPI (Blue). Yellow arrows denote mitotic chromosomes with associated TRIP11 specks. Note the correspondence of staining of the Golgi apparatus, comparing GIANTIN (middle row) with TRIP11 (top row). Also note the almost complete absence of fine green specks in granulocytic sh1 cells, compared to their presence in granulocytic S4 cells. Magnification bar: 10 μm.

### Gene Ontology Analysis-Acetyltransferase and Methyltransferase Complexes

Based upon assigned gene functions in GeneCards (genecards.org), the “only-sh1-down” gene datasets in (GO:0034708) “Methyltransferase Complex” and in (GO:1902493) “Acetyltransferase Complex”, revealed a number of genes with apparently contradictory roles. For example, in the Methyltransferase Complex, the genes KDM5B and KDM5C are described as lysine demethylases; PRDM4 and PRDM10 are methyltransferases. Yet, all 4 genes (as well as others with similar functions) exhibit transcript downregulation. Within the Acetyltransferase Complex, the genes KAT8 and JADE2 are histone lysine acetyltransferases; but they are also downregulated. The overall impression is that sh1 cells do not exhibit much change in histone lysine methylation or acetylation, comparing undifferentiated with RA-differentiated cell forms.

This interpretation is supported by immunostaining experiments (Figure 10) employing antibodies specific for methylated or acetylated histones within interphase nuclear chromatin. As examples, Figure 10a compares H3K9me2 between S4 and sh1 cells, each cell line without or with RA treatment. In all four cases, anti-H3K9me2 appears to be most concentrated close to the interphase NE and less in the nuclear center. Also, anti-H3K9me2 does not appear to stain ELCS (yellow arrowhead) or the perinucleolar chromatin region (thin yellow arrows). Figure 10b reveals that anti-H3K9me3 appears to stain ELCS and sites scattered around the granulocyte nuclear lobes. Figure 10c presents staining by anti-H3K4,8,12,16 multi-acetylation, comparing S4 and sh1 RA treated granulocytes. The multi-acetylation antibody does not appear to stain ELCS in the S4 RA treated cells; but stains in a spotty manner throughout the S4 and sh1interphase chromatin. This observation, combined with the indication of staining by anti-H3K9me3 is consistent with the idea that ELCS chromatin is transcriptionally inactive. As mentioned earlier, cryo-electron microscopic tomography demonstrates that ELCS chromatin is structured with two layers of ∼30 nm diameter chromatin fibers in a “crisscross” pattern, with the fibers running parallel to the adjacent INM (Xu et al., 2021).

**Figure 10.**
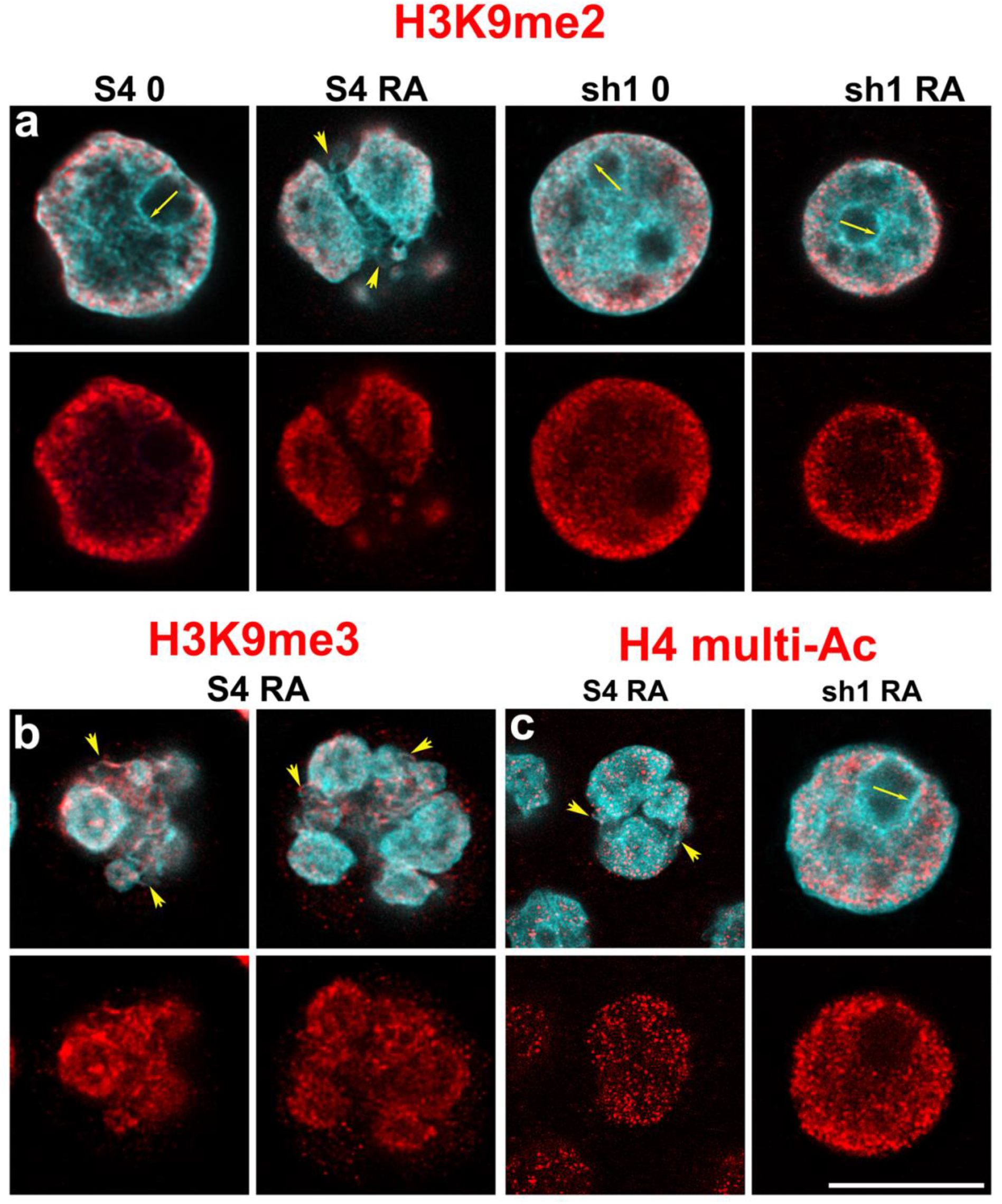
Immunostaining HL-60/S4 and sh1 cells for histone methylation and acetylation. **a**) Anti-H3K9me2 staining of four cell states: S4 0; S4 RA; sh1 0; sh1 RA. **b**) Anti-H3K9me3 staining S4 RA (two examples) to demonstrate staining of ELCS. **c**) Anti-H3K4,8,12,16 multi-acetylation antibody staining two cell states: S4 RA and sh1 RA. The bottom row in each set of images displays the antibody-only stained in red; the top row displays a merge of DNA staining (cyan) and antibody staining (red). Note that anti-H3K9me2 does not stain ELCS (yellow arrowheads) in (**a**) S4 RA, whereas anti-H3K9me3 appears to stain ELCS in (**b**) S4 RA. There is possible weak staining of ELCS by anti-H3K4,8,12,16 multi-acetylation antibody in (**c**). Perinucleolar chromatin regions (thin yellow arrows) appear not to be stained employing anti-H3K9me2 or anti-H3K4,8,12,16 multi-acetylation antibody. Magnification bar: 10 μm.

### Gene Ontology Analysis-Centromere Effects

Examination of the chromosome-related downregulated gene sets (see GO terms, Table 2) revealed a diversity of consequences in sh1 cells, compared to S4 cells. The most obvious consequence appears to be at the centromere/kinetochore region of chromosomes. Of the 16 genes identified in (GO:0000793) “Condensed Chromosome”, 7 “only-sh1-down” genes play roles in the normal structure and function of the centromere/kinetochore. In addition, 3 other “centromere/kinetochore” genes are downregulated in sh1 cells, but not “only-sh1-down”. These 10 genes are presented (Figure 11) as Log_2_FC downregulated sh1 transcripts compared to the transcript levels in S4 cells, following treatment with retinoic acid (the plots are based upon Table S2 “pairwise”). It is important to note that sh1 and S4 grow and divide in their respective media, and <u>both</u> gradually cease dividing when treated with RA. It is conceivable that the cellular differences in transcription levels reflect a differential dismantling of the centromere/kinetochore structure as the cells cease mitosis. The transcript levels of CENPA clearly decrease in both RA-treated sh1 and S4 cells. The other genes decrease with less magnitude and less statistical significance. The reason why sh1 cells exhibit more dramatic changes is not clear. Perhaps, the sh1 cells cease mitosis more rapidly than S4 cells.

**Figure 11.**
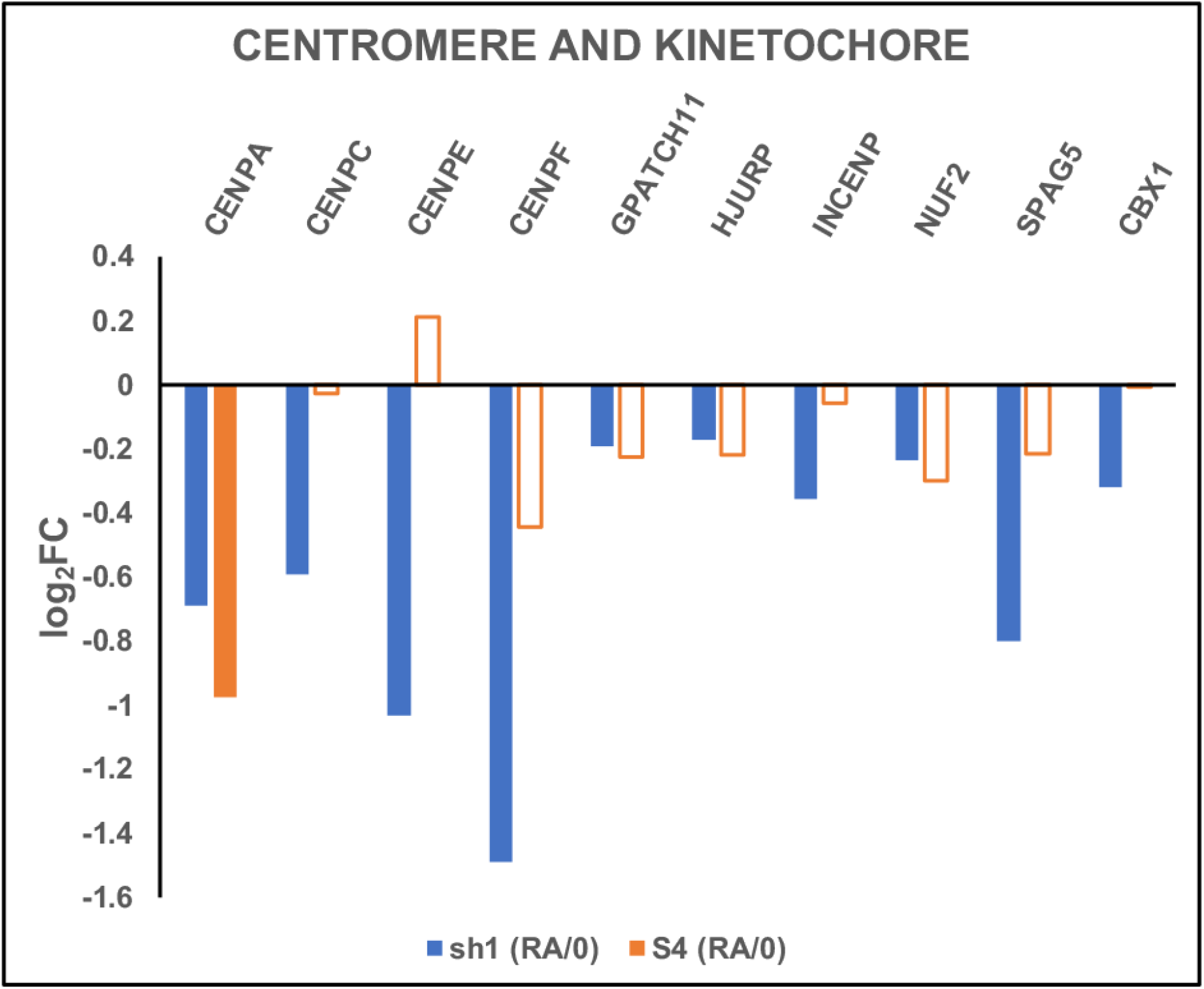
Downregulation of centromere/kinetochore protein transcripts resulting from RA treatment, comparing the responses of sh1 to S4 cells. Closed bars: PPDE>0.95 (significant data). Open bars: PPDE<0.95 (not significant). Plots are derived from data in Table S2 “pairwise”. HGNC Gene codes are displayed above the relative transcript level bars.

### Gene Ontology Analysis-DNA Damage and Repair

Based upon assigned gene functions in GeneCards, the “only-sh1-down” gene datasets in (GO:0000793) “Condensed Chromosome” and (GO:0000151) “Chromosomal Region”, revealed a number of genes with roles in DNA damage repair (Figure 12). Downregulation of these genes in sh1 cells supports the speculation that sh1 cells may be more sensitive to DNA damage than the S4 cell type.

**Figure 12.**
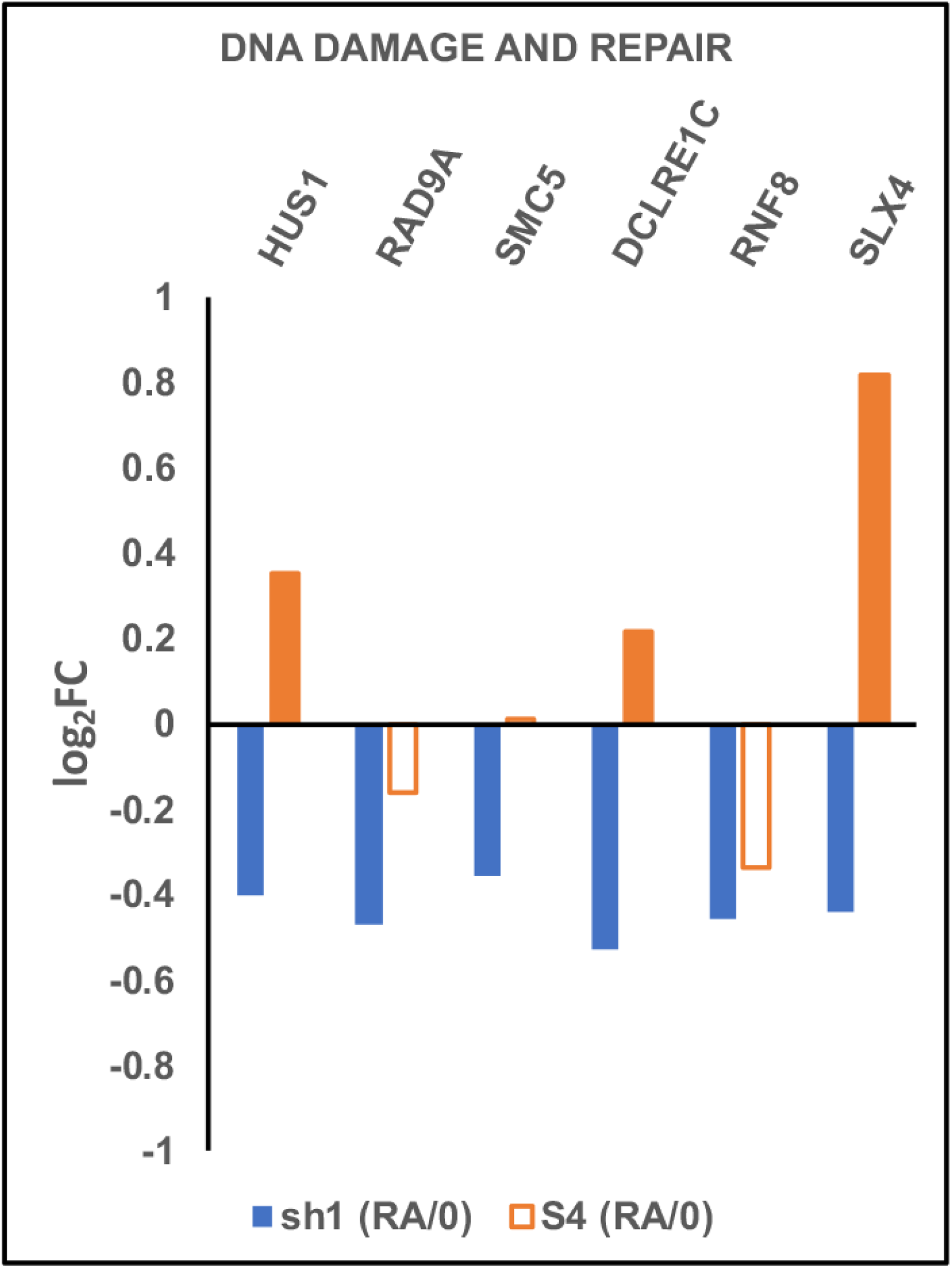
Downregulation of DNA damage and repair protein transcripts resulting from RA treatment, comparing the responses of sh1 to S4 cells. Closed bars: PPDE>0.95 (significant data). Open bars: PPDE<0.95 (not significant). Plots are derived from data in Table S2 “pairwise”. HGNC Gene codes are displayed above the relative transcript level bars.

### Gene Ontology Analysis-Ribosomes

The only GO term that is statistically significant (i.e., FDR ≤0.05) in the “Only-sh1-up” gene list is: GO:0005840 “Ribosome”. This GO term contains 19 genes, mostly structural ribosomal proteins for the large and small ribosomal subunits; also, a few genes assisting mRNA translation. Figure 13 clearly demonstrates that sh1 cells show significant ribosome gene upregulation when treated with RA; whereas there is considerable downregulation of ribosome transcripts when S4 cells are treated with RA. This observation suggests that RA treated sh1 cells are performing *increased* ribosome protein synthesis compared to RA treated S4 cells, which appear to have *decreased* protein synthesis. Turnover of the ribosomal proteins in S4 and sh1 cells should be studied by existing methods (Mathis et al., 2017). Despite the observation that ribosomal protein transcripts are increased in LBR knockdown cells (HL-60/sh1) that are differentiated into granulocytes with RA, the transcript levels for RNA polymerases (pol) I, II and III protein subunits are largely downregulated after RA treatment in <u>both</u> sh1 and S4 cells (Figure 14). This implies a reduction in ribosomal structural RNA (i.e., the pol I 45S pre-rRNA, ultimately processed into 28S, 18S and 5.8S RNA and the pol III tRNA and 5S RNA) in <u>both</u> sh1 and S4 cells. The downregulation of RNA pol II protein subunit genes could lead to reduction of various mRNAs, but is not evident with regard to ribosomal protein transcripts (Figure 13).

**Figure 13.**
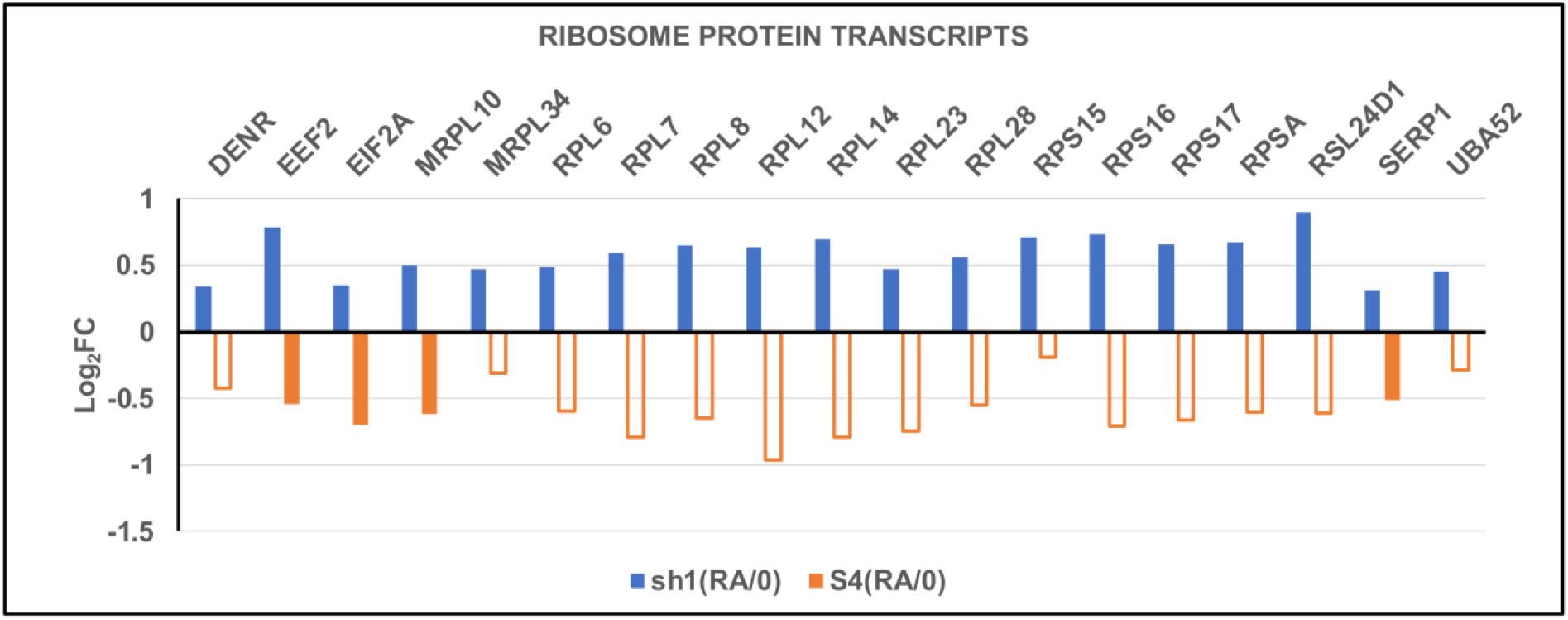
Upregulation of ribosomal protein transcripts resulting from RA treatment of sh1 cells, compared to the downregulation responses of RA treatment of S4 cells. Closed bars: PPDE>0.95 (significant data). Open bars: PPDE<0.95 (not significant). Plots are derived from data in Table S2 “pairwise”. HGNC Gene codes are displayed above the relative transcript level bars.

**Figure 14.**
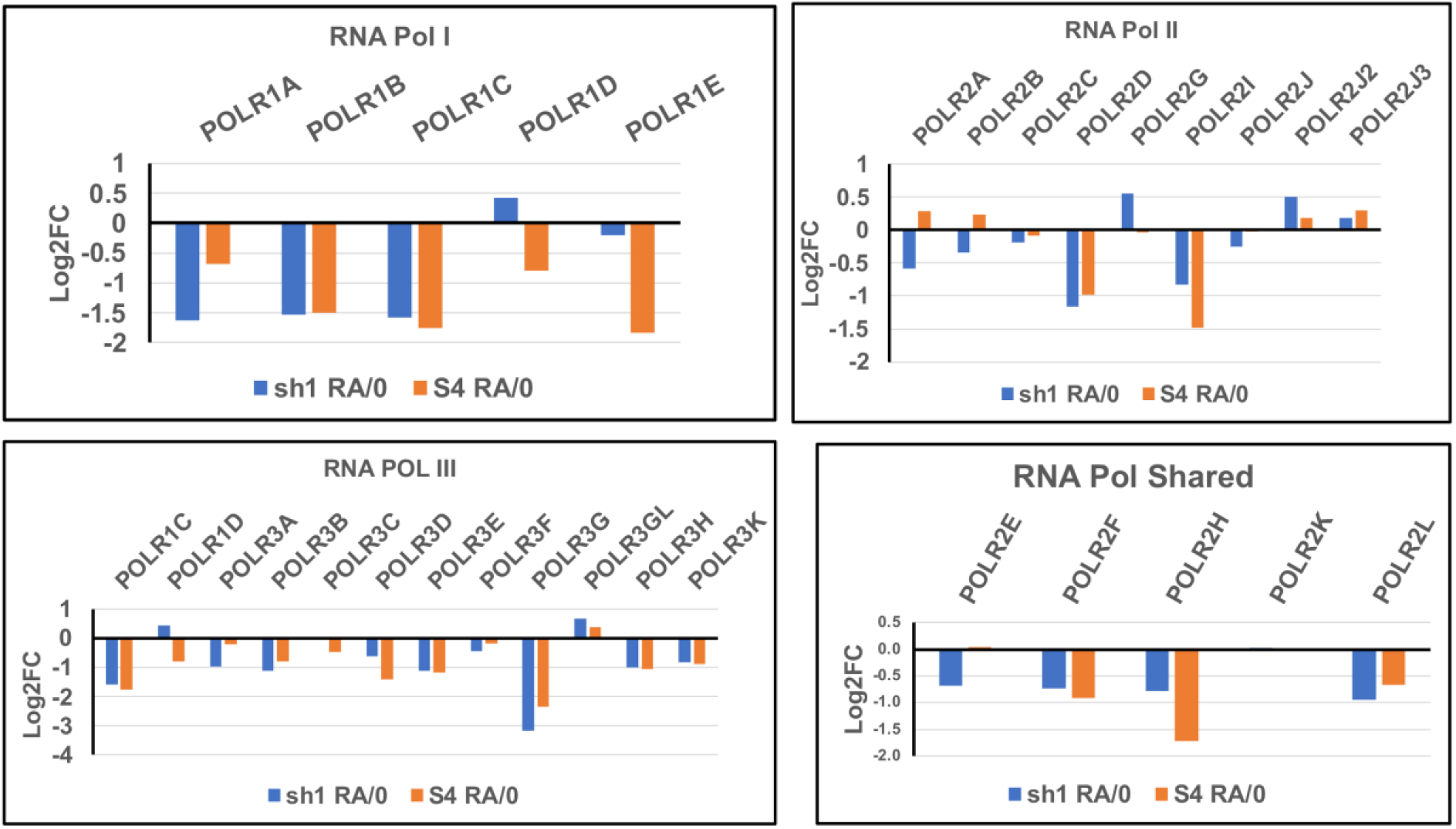
Relative transcript levels for protein subunits of RNA polymerases I, II and III, comparing HL-60/S4 and HL-60/sh1 cells, each cell type retinoic acid treated versus undifferentiated (RA/0). Polymerase-specific subunit transcripts are displayed in individual panels, with shared subunit transcripts in a separate panel. See Table S2 “pairwise” for complete data. Y axis: Log2FC, log2 of the ratio of transcript levels RA versus 0. Open bars signify that the change in transcript level is not significant (PPDE<0.95). Solid bars signify that the change in transcript level is significant (PPDE>0.95). HGNC Gene codes are displayed above the relative transcript level bars.

### LBR and the Nucleolus

Ribosomal RNA synthesis and ribosome assembly with structural rRNA and proteins occurs principally within the nucleolus (Bizhanova and Kaufman, 2021; Panov et al., 2021; Potapova and Gerton, 2019). The consensus model of an interphase nucleolus performing ribosome biosynthesis involves sequential activity in the following regions: 1) rRNA synthesis in nucleolar fibrillar centers (FC) containing the rDNA repeats; 2) processing rRNA in the dense fibrillar component (DFC), adjacent to the FCs; 3) addition of ribosomal proteins in the granular component (GC), surrounding the DFCs; 4) passage of the 40 and 60S pre-ribosomal subunits through the perinucleolar heterochromatin “shell” around the GC; 5) final assembly of ribosomes in the cytoplasm (Kang et al., 2021).

For many years, in numerous microscopy experiments, we have observed in HL-60/S4 cells (0 and RA) that, in a high percentage of cells, the interphase nucleoli exhibit close proximity or contact with the nuclear envelope. By contrast, in HL-60/sh1 cells (0 and RA) interphase nucleoli often appear to be distant from the nuclear envelope, sometimes in the “center” of the nucleus. Examples of these general observations can be seen in Figures 2, 10, 15-18.

**Figure 15.**
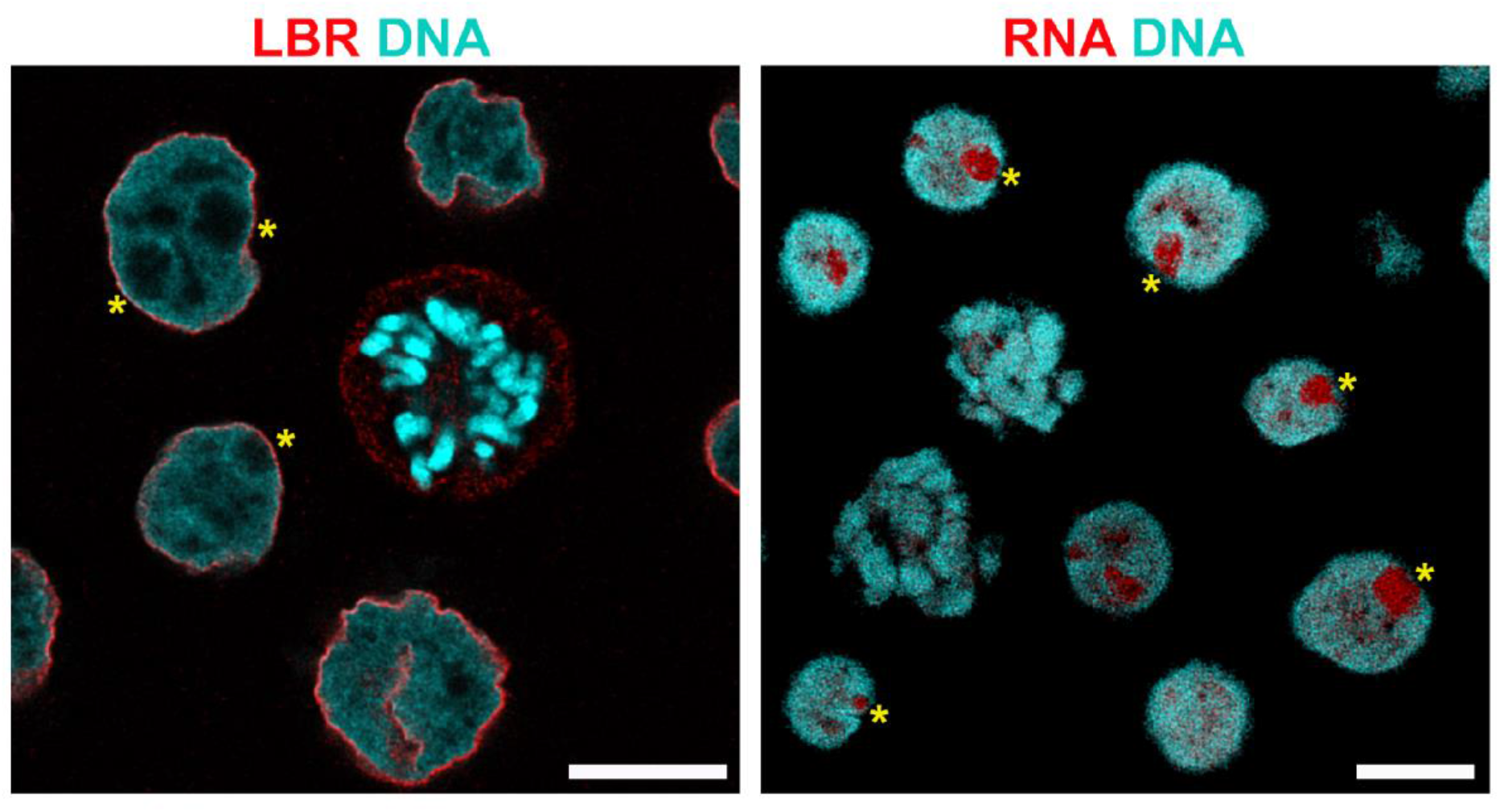
Undifferentiated HL-60/S4 cells. Left frame: immunostained for LBR (red) and DAPI stained DNA (cyan). Yellow asterisks denote presumptive nucleoli adjacent to the interphase nuclear envelope. In the mitotic cell, LBR is dispersed following nuclear envelope breakdown. Right frame: RNA specific staining with SYTO RNASelect (ThermoFisher Scientific). Yellow asterisks denote presumptive nucleoli (red) adjacent to the periphery of the interphase nuclear chromatin staining (cyan). Magnification bar: 10 μm.

**Figure 16.**
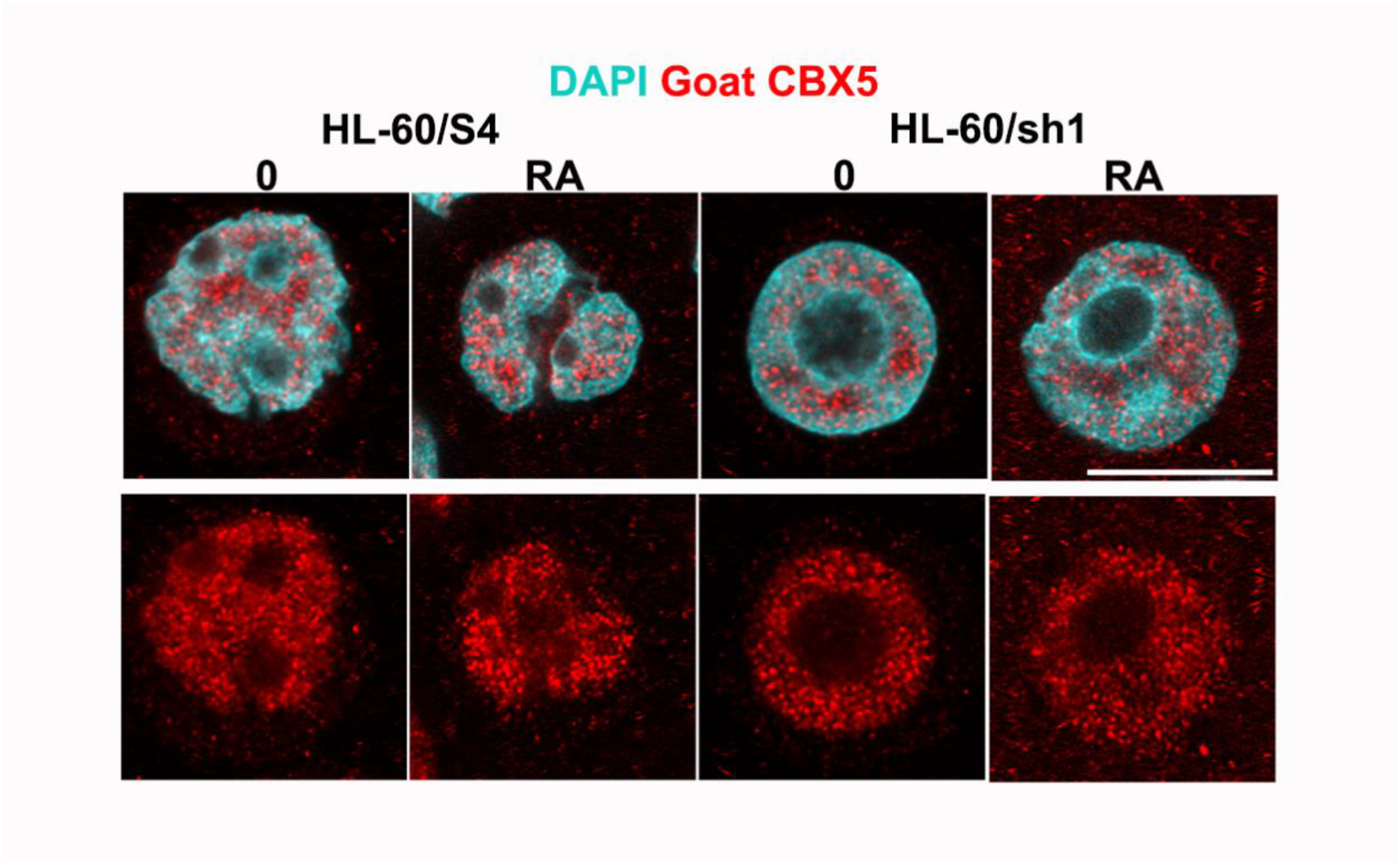
Immunostaining of HL-60/S4 and HL-60/sh1cells with goat anti-CBX5 (HP1a) (red) and DNA (DAPI, cyan). Each cell-type is displayed both undifferentiated (0) and retinoic acid differentiated (RA). Note that the presumptive nucleoli and perinucleolar chromatin are devoid of CBX5. Some nucleoli are adjacent to the nuclear envelope (esp., HL-60/S4, 0 and RA), while some are more central in the nuclei (HL-60/sh1, 0 and RA). Magnification bar: 10 μm.

**Figure 17.**
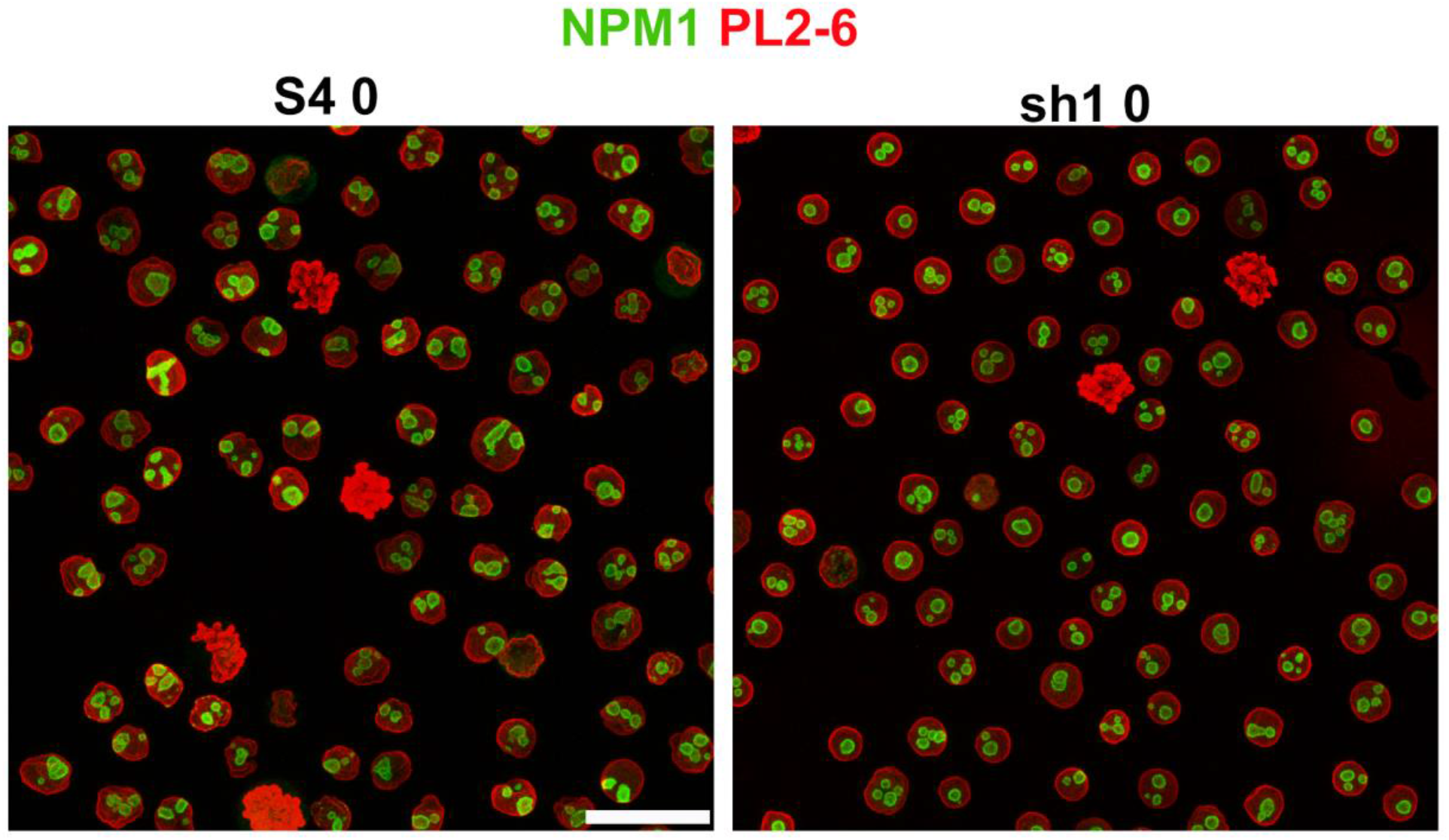
Immunostaining survey of the number of nucleoli/nucleus in undifferentiated HL-60/S4 (left frame) and HL-60/sh1 cells (right frame). The cells were stained with anti-NPM1 (green) and anti-epichromatin (red). Anti-NPM1 stains at the periphery of each nucleolus. Both image frames are Maximum Intensity Projections (MIP). Mitotic chromosomes are also stained at their periphery with anti-epichromatin. However, after MIP each cluster of mitotic chromosomes appears completely red. Superimposing a “grid” (not shown) over each frame, we examined each nucleus and categorized it as containing 1, 2 or >2 nucleoli. Tiny nucleolar fragments were not included in the counting, nor were mitotic included. The number of S4 0 cells counted was 83; sh1 0 cells, 86. Magnification bar: 10 μm.

**Figure 18.**
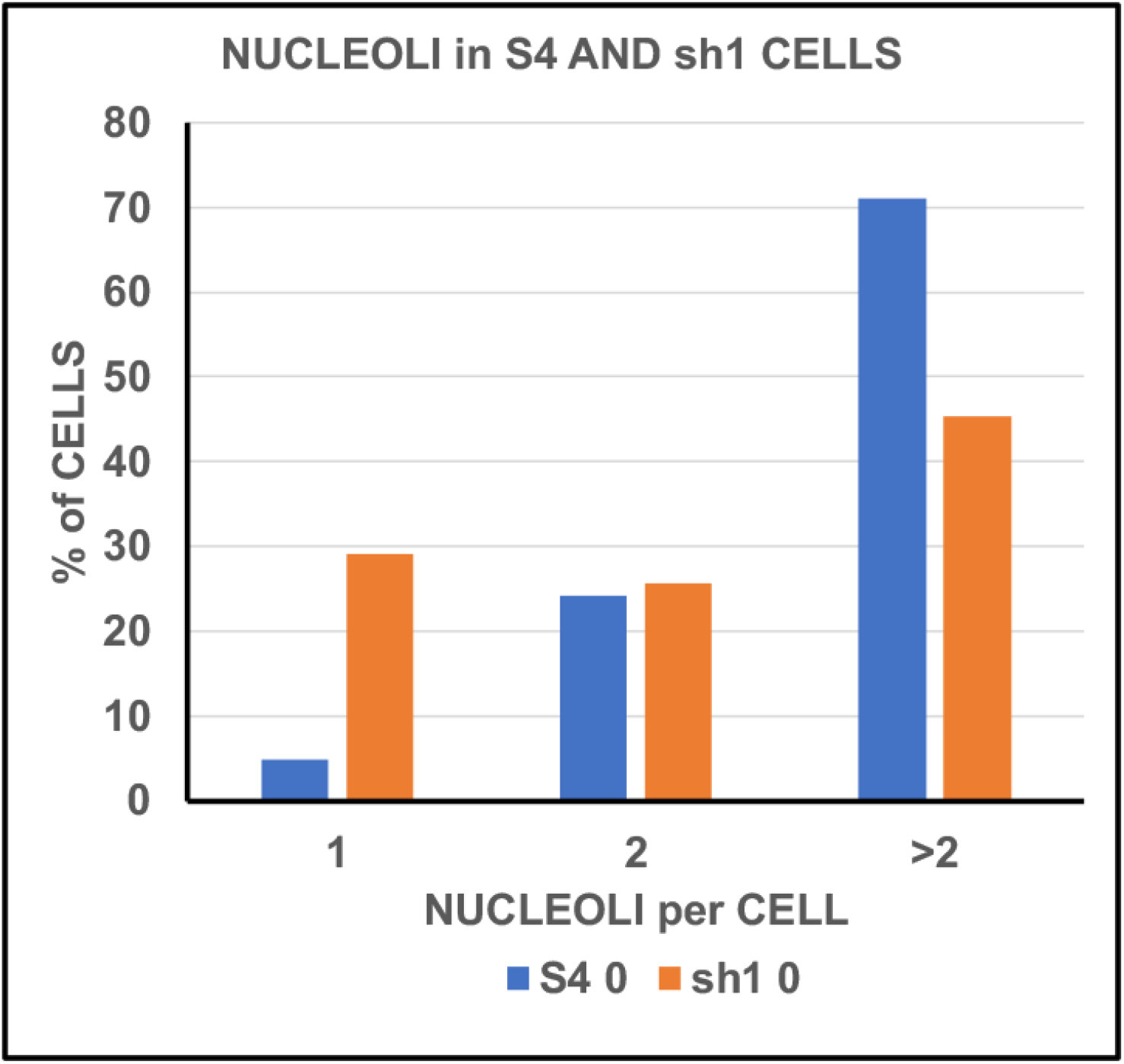
The percentage of undifferentiated HL-60/S4 and of HL-60/sh1 cell nuclei exhibiting 1, 2 or >2 nucleoli/nucleus. About 30% of the HL-60/sh1 cells exhibit one nucleolus/nucleus, generally separated from the nuclear envelope. By contrast, ∼70% of the HL-60/S4 cells exhibit >2 nucleoli/nucleus, with many being close to the nuclear envelope.

These repeated observations about nucleolar positions within the interphase nuclei prompted us to develop a more quantitative overview. For this purpose, we examined, categorized and counted the immunostained images of undifferentiated HL-60/S4 and HL-60/sh1 cells (Figure 17), utilizing anti-nucleophosmin (NPM1) to visualize nucleoli (green) and anti-epichromatin (using mAb PL2-6) to visualize the nucleosome acidic patches (red) at the nuclear chromatin periphery (Gould et al., 2017; Zhou et al., 2019). The final count of the number of nucleoli/nucleus in each established category (i.e., 1, 2 or >2 nucleoli/nucleus) are presented in Figure 18. This count supports the conclusion that ∼70% of interphase S4 nuclei have >2 nucleoli/nucleus, whereas a minority (∼45%) of interphase sh1 nuclei have >2 nucleoli/nucleus. About 30% of interphase sh1 nuclei have only one nucleolus/nucleus.

## Discussion

The present study attempts to discover cellular functional consequences of LBR deficiency in the myeloid leukemia cell line HL-60/S4, beyond the structural microscopic observations (Olins et al., 2010a). We have previously shown that LBR “knockdown” in HL-60/S4 cells (the “sh1” cell line) blocked RA-treated granulocytic forms from developing nuclear lobulation and Envelope-Limited Chromatin Sheets (ELCS) (Olins et al., 1998; Olins et al., 2010b; Olins and Olins, 2009). To uncover functional information, in the present study we have analyzed the polyA mRNA transcriptomes of sh1, S4 and gfp (a vector “control”) cell lines. We “curated” the sh1 transcriptome into two mapped gene lists; i.e., “only-sh1-down” and “only-sh1-up”, requiring that sh1 transcript levels following RA exposure change in the **opposite** direction, compared to **both** S4 **and** gfp cell types. These curated gene lists are much smaller than the total mapped gene transcripts (e.g., 638 “only-sh1-down” and 696 “only-sh1-up”, versus 16,822 total genes; see Table 1). These curated gene lists were then subjected to Gene Ontology (GO) analysis. Identified statistically significant GO terms (FDR≤0.05) contained subsets of these curated gene lists, designated by us as “gene datasets”. These datasets were further scrutinized. Gene functions were derived from GeneCards (www.genecards.com), forming the basis of our analyses and speculations.

We had expected that the top “only-sh1-down” GO term would relate to chromatin structure; but this was not what we observed (Table 2). The most significant GO term in “only-sh1-down” is (GO:0005801) “Cis-Golgi Network”. The definition of this GO term is: “The network of interconnected tubular and cisternal structures located at the convex side of the Golgi apparatus, which abuts the endoplasmic reticulum” **. The implication of this GO term “only-sh1-down” is that the “network” may not be functioning efficiently (e.g., nascent proteins may not be moving into the Golgi stacks, and protein modification and secretion may be faulty) in these LBR-knockdown cells. A search of scientific publications, queried as “LBR and Golgi”, turned up only one primary research paper (Wehrle et al., 2018), also cited in a review (Lowe, 2019). The authors of this primary research paper describe a lethal human skeletal disease (Chondrodysplasia achondrogenesis 1A, “ACG1A”) caused by a mutation in the TRIP11 gene (which codes for the Golgin protein GMAP-210). They point out that a similar phenotype is seen in a homozygous human LBR mutation; a syndrome named “Greenberg Dysplasia” (Greenberg et al., 1988). The authors suggest that there is a common disruption of Golgi apparatus architecture resulting from the TRIP11 or from the LBR mutation, which adversely affects the secretory function.

This opens the question of a possible role of LBR in the control of golgin gene expression. One might speculate that LBR has a “direct” gene regulatory effect upon TRIP11 gene transcription (located at Chr 14q32.12). Numerous golgin genes are scattered throughout the human chromosome set, with the exception of the Golgin A8 family, which is located at Chromosome 15q13 and 14. Our transcriptome analysis indicates that sh1 cells (+/- RA) exhibit considerable downregulation of TRIP11 gene expression **and** many other Golgi protein genes, compared to their expression in S4 cells (Figure 8). This argues against a direct “single” effect of LBR knockdown, confined to the TRIP11 gene.

A second speculation could be that the LBR deficiency results in local reduction of sterols within Golgi cisternae and vesicle membranes. A recent study (Young et al., 2021) on a mutant mouse line with an LBR N-terminal tail (236 bp) truncation indicated defects in GO terms involving “endomembranes” of the Nuclear Envelope (NE) and the ER. It is important to mention that comparative immunostaining of sh1 cells (+/- RA), using the “Golgi marker” anti-Giantin (GOLGB1) does not indicate any major changes in Golgi shape and stain intensity (Figure 9). However, Figure 9 <u>does</u> reveal that putative Golgi vesicles stained with anti-TRIP11 are present in RA-treated S4 cells; but less apparent is RA-treated sh1 cells. This observation would be in agreement with the interpretation that Golgi vesicle architecture is perturbed (Wehrle et al., 2018). In addition, we have immunostaining evidence that LBR does <u>not</u> localize within the Golgi apparatus (Figure 19). We suggest that the endomembranes of NE and ER vesicles in sh1 cells, traveling to the cis-Golgi network, have a deficiency in sterols due to LBR knockdown, which may affect the Golgi membrane structure and function.

**Figure 19.**
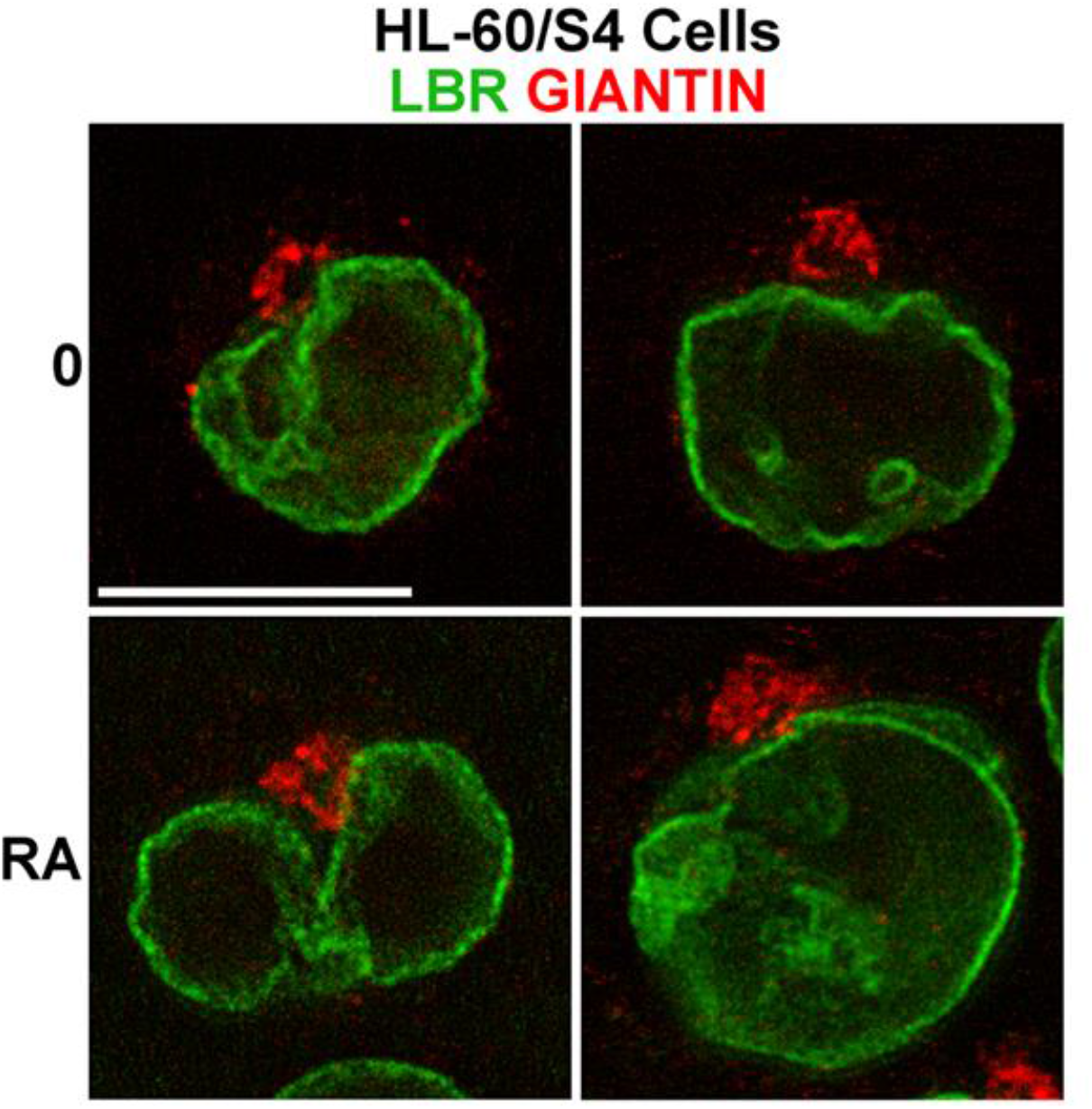
Immunostaining of HL-60/S4 cells with anti-LBR (green) and anti-Giantin (red). Top row, undifferentiated S4 cells; bottom row, granulocytic S4 cells. Note the apparent absence of LBR in the Golgi apparatus. Magnification bar: 10 μm.

A third speculation is that the N-terminal tail with the Tudor Domain and HP1 binding motif are associated with many different heterochromatic regions containing a broad diversity of repressed genes. When there is a knockdown of LBR, as in sh1 cells, many seemingly unrelated cell functions can be perturbed, in addition to the more understandable functions of NE membrane growth and nuclear shape changes (Olins and Olins, 2009). Heterochromatin rearrangements in sh1 cells may result in downregulation of genes with no obvious relationship to LBR (e.g., “Centromere effects” or “DNA damage and repair”). These hypothesized rearrangements of heterochromatin in sh1 cells, compared to S4 cells could probably be detected and genetically evaluated employing Hi-C to determine perturbations in chromatin compartments (Lieberman-Aiden et al., 2009). It would also be important to locate LBR bound to chromatin, identifying the corresponding genomic locations, by employing variations of ChIP-seq technology.

The discovery that the most significant “only-sh1-up” GO term is “Ribosome” was also a surprise. As a consequence, LBR knockdown results in upregulation of ribosome protein transcripts in RA-treated HL-60/sh1 cells, but not in RA-treated HL-60/S4 cells (Figure 13). Combined with downregulation of RNA polymerase subunit transcript levels (Figure 14), generates a “true” conundrum. Regulation of ribosome protein genes has been recently reviewed (Kofler et al., 2020; Petibon et al., 2021). This complex topic is beyond the scope of the present article. Indeed, to quote one of the above reviews: “This whole maturation cascade is driven by about 250 different assembly factors each required at a distinct stage.” (Kofler et al., 2020).

As described in the **Results** and demonstrated in Figures 2, 10 and 15-18, the presence or absence (i.e., knockdown) of LBR affects the number of nucleoli/nucleus and their proximity to the nuclear envelope. Normally, LBR is not located in interphase perinucleolar heterochromatin. We speculate that during interphase of HL-60/S4 cells, LBR acts as a “bridge” between the nuclear envelope and perinucleolar heterochromatin. In HL-60/sh1 cells, this “bridge” is much reduced, permitting the nucleolus with its heterochromatin “shell” to acquire a more central position within the nucleus. Furthermore, the nucleolus is regarded as a prime example of a non-membranous multiphase nuclear “condensate”, stabilized by liquid-liquid phase separation (Lafontaine et al., 2021; Yoneda et al., 2021). Multiple nucleoli can be coalesced or dispersed by changes in the local biophysical environment. Our perspective is that the presence of sufficient LBR in the nuclear envelope will “bridge” to the perinucleolar heterochromatin, stabilizing the multiple nucleolar condensates, whereas, LBR deficiency “liberates” the perinucleolar heterochromatin creating a more permissive nuclear environment for coalescence of the multiple nucleoli. We suspect, but cannot prove, that these changes in the nuclear and nucleolar biophysical environment are, in part, responsible for the transcriptional changes inherent in the “only-sh1-up” GO term “Ribosome”.

It would be useful to extend the present study of LBR knockdown in human HL-60/S4 cells to the mouse LBR “Ichthyosis” (*ic*) mutation (Hoffmann et al., 2002; Shultz et al., 2003). We would like to determine whether mouse TRIP11 is downregulated in parallel to the LBR downregulation, as observed in the human HL-60/S4 and HL-60/sh1 cell lines. We would like to see whether nucleolar position, number and transcription are also affected by LBR knockdown. A progenitor mouse EML (erythroid, myeloid & lymphoid potential) derived promyelocyte with a homozygous *ic* mutation can be differentiated to granulocyte form following RA treatment, and displays a “kidney-shaped” nucleus, instead of the normal “ring-shaped” nucleus (Gaines et al., 2008). For a comprehensive study to commence, mRNA transcriptomes will need to be obtained for the undifferentiated and RA-differentiated mouse normal and Ichthyosis promyelocytes (Zwerger et al., 2008). It would also be useful if transcription data becomes available for the earlier mentioned mouse cell line with an N-terminal truncation of LBR that exhibits PHA (Young et al., 2021).

## Conclusion

Combining immunostaining microscopy and mRNA transcriptome analysis, we have explored additional functions (i.e., beyond those published) for Lamin B Receptor (LBR) in human myeloid HL-60/S4 cells and in “LBR knockdown” HL-60/sh1 cells, in both the undifferentiated (0) and granulocytic (Retinoic Acid treated, RA) state. Gene Ontology (GO) analysis of “only-sh1-down” (i.e., decreased transcription) and “only-sh1-up” (i.e., increased transcription), compared to control cell lines HL-60/S4 and HL-60/gfp, yielded surprising results. The “top” GO term for “only-sh1-down” was “Cis-Golgi-Network”; the top for “only-sh1-up” was “Ribosome”. Speculations are presented for how an LBR knockdown can yield these altered cellular physiological states.

## Acknowledgments

The authors express their gratitude to the Woods Hole Marine Biology Laboratory. DEO and ALO also thank the School of Pharmacy, University of New England for space and support for the past 11 years. In addition, DEO and ALO express their gratitude Dr. Igor Prudovsky at the MaineHealth Institute for Research for inviting them to join his research group.

**The definition of “Cis-Golgi Network” according to (QuickGO:Term GO:0005801) “The network of interconnected tubular and cisternal structures located at the convex side of the Golgi apparatus, which abuts the endoplasmic reticulum. The CGN is not considered part of the Golgi apparatus but is a separate organelle.

## Notes

### Competing Interest Statement

The authors have declared no competing interest.

